# « Fluorescent Protein Photobleaching: From molecular processes to spectromicroscopy »

**DOI:** 10.64898/2026.03.31.715555

**Authors:** Théo Beguin, Kaixin Wang, Yasmina Bousmah, Ninette Abou Mrad, Frederic Halgand, Hélène Pasquier, Marie Erard

## Abstract

Fluorescent proteins (FPs) are essential tools for biological imaging but are limited by photobleaching, a light-induced loss of fluorescence intensity that reduces spatial and temporal resolution. Despite extensive use, the molecular mechanisms underlying FP photobleaching remain poorly understood due to the diversity of FPs and the complexity of their photochemistry. Existing approaches either monitor fluorescence decay in live cells, reflecting imaging conditions but lacking molecular detail, or rely on in vitro spectroscopy of purified proteins, providing mechanistic insight but often limited to individual FPs.

We introduce a quantitative workflow bridging these approaches by combining live-cell measurements with in vitro spectroscopy. In vitro measurements are performed on a dedicated setup that simultaneously monitors absorption, emission, and fluorescence decay during photobleaching. Applied to six FPs spanning different chromophores, emission ranges and sequences, this approach reveals that photobleaching strongly depends on FP. It involves multiple chemical pathways, including oxidation, dimerization, and backbone cleavage. Spectroscopic analysis uncovers a heterogeneous ensemble of photoproducts with distinct photophysical properties that can remain optically active during irradiation, including shortened fluorescence lifetimes or altered absorption spectra.

These findings demonstrate that FP photobleaching cannot be described as a simple ON–OFF process but involves complex transformations affecting both fluorescence intensity and lifetime. Such transformations can introduce significant biases in quantitative imaging, particularly in advanced techniques such as FLIM and FRET. Finally, we introduce quantitative indicators enabling robust comparison of FP photostability across experimental conditions. This framework provides a comprehensive approach for understanding and quantifying photobleaching and its implications for fluorescence imaging.

## Introduction

Fluorescent proteins (FPs) are genetically encoded probes that have become essential tools in biological imaging, enabling the investigation of chemical and biochemical processes at the molecular level and in living cells^1,2^. FPs have been identified in numerous marine organisms over the past 30 years^3^. They share a common structural motif: a β-barrel architecture, typically composed of 11 β-strands that form a cylindrical shell enclosing a central α-helix (Scheme 1). Within this helix, the chromophore is generated through the autocatalytic cyclization of three amino acids, usually involving an aromatic residue—most often a tyrosine—together with a glycine in the third position (X–Tyr–Gly). The β-barrel structure plays a critical protective role by shielding the chromophore from solvent-induced quenching while providing a rigid environment supported by an extended hydrogen-bond network, which is necessary for efficient fluorescence^4^. Variations in the residues surrounding the chromophore, as well as engineered mutations, can modulate properties such as folding rate, maturation efficiency, excitation/emission wavelengths, brightness, pH sensitivity, and photostability.

Accurate monitoring of biological events is often limited by FP photobleaching—a progressive loss of fluorescence intensity upon light excitation—which compromises both the temporal and spatial resolution of imaging experiments^5,6^. The mechanisms underlying FP photobleaching, as well as the detailed characterization of the resulting modifications to their photophysical properties, remain poorly understood. This is largely due to the wide diversity of fluorescent proteins, which can exhibit different light-induced reactivities depending on both the chemical nature of their chromophore and their overall protein sequence^7^. As a result, photobleaching studies are typically constrained to two main approaches. The first focuses on monitoring the rate of decrease in fluorescence intensity in cells expressing FPs during microscope irradiation. This strategy reflects the conditions of biological imaging and is widely used to compare the photostability of numerous FPs across a broad range of light intensities^5^. The second approach involves detailed characterization of photochemical reactions within FPs by combining, in vitro, spectroscopic and structural techniques. It is typically limited to a specific purified protein. This approach has allowed the identification of various chemical processes, such as the photocleavage in EGFP^8^ and the decarboxylation of Glu222 in EYFP^9^, Glu212 in IrisFP^10^ or Glu215 in DsRed^11^. To date, the most comprehensive study of this type was performed on a crystal of IrisFP, for which a complete photobleaching mechanism was proposed, including the influence of experimental conditions such as excitation power density and the presence of molecular oxygen^10^.

Here, we propose a workflow to analyse the photoproducts generated during FP photobleaching and to assess how these species affect both intensity- and lifetime-based imaging applications. Monitoring the photoinduced decay of fluorescence intensity under a microscope in live cells prevents detailed spectroscopic characterization of the underlying photochemical processes. To overcome the limitations of previous studies, we developed a dedicated spectroscopy setup, called BEAM (Bleaching Emission and Absorption Monitoring), which allows real-time monitoring of absorption and emission spectra together with fluorescence intensity decay during the photobleaching of purified FPs ^12^.

This workflow was applied to six FPs derived from either *Aequorea victoria* or *Discosoma* species, which share a similar overall structure but exhibit completely distinct primary sequences (Scheme 1, see SI, part A). The green FP EGFP, the yellow FPs EYFP and Citrine (EYFP Q69M), and the red FP mCherry contain tyrosine-based chromophores, whereas the cyan FPs Aquamarine and mTurquoise contain tryptophan-based chromophores. This selection also enables the exploration of the role of specific residues, as EYFP and Citrine differ by only one mutation, and Aquamarine and mTurquoise by four. In addition, these FPs were chosen because of their widespread use in fluorescence microscopy, particularly in Förster Resonance Energy Transfer (FRET) experiments, where CFP/YFP and GFP/RFP are among the most commonly employed FRET pairs.

**Scheme 1:**
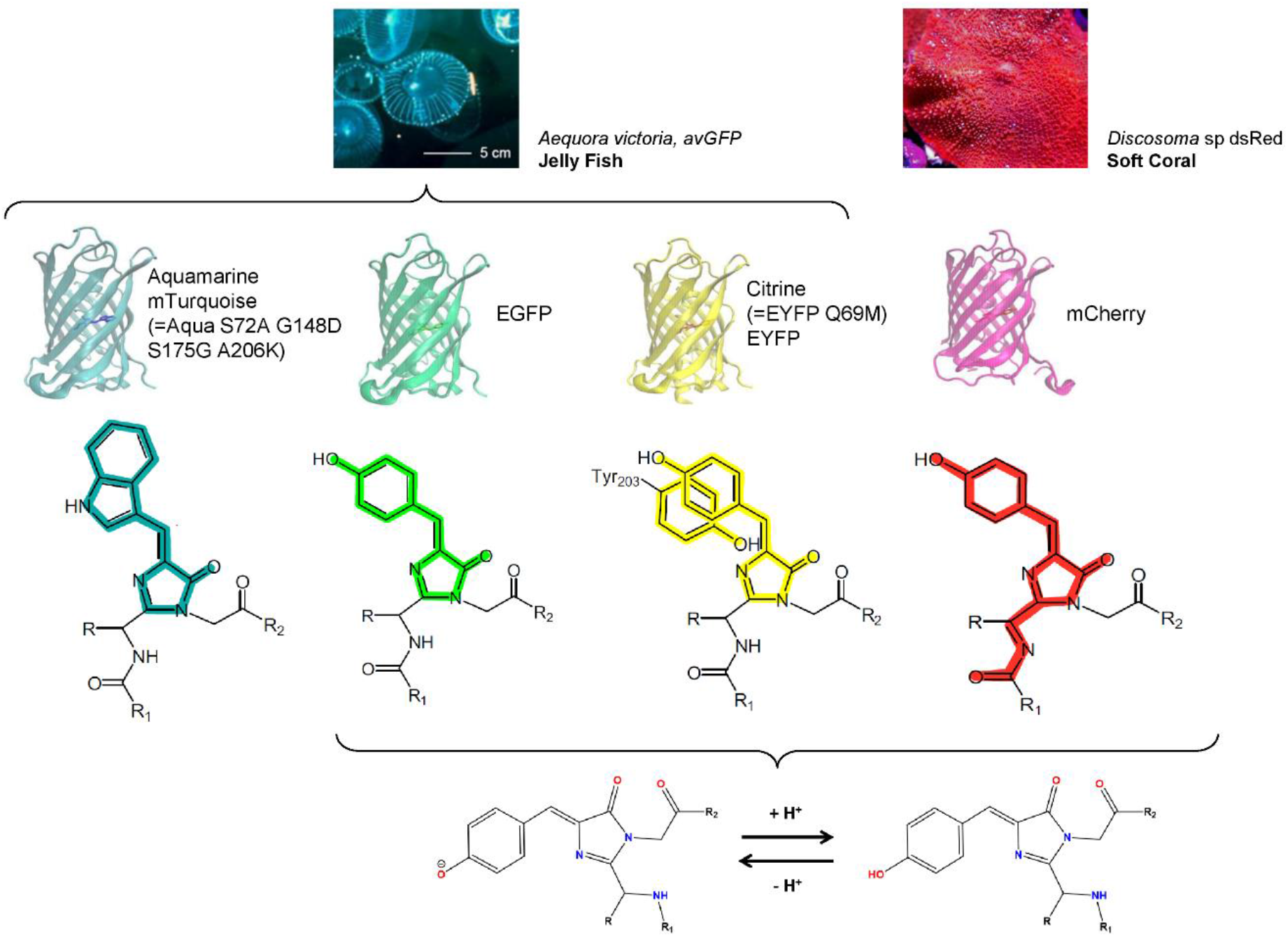
The photobleaching of six FPs is investigated, spanning four emission wavelength ranges and types of chromophores, all derived from two marine organisms. At the bottom, the acid–base equilibrium of the tyrosine-based chromophores is shown. In Citrine, EYFP, EGFP, and mCherry, only the anionic form of the chromophore is fluorescent.

To ensure that in vitro observations are representative of those obtained in live-cell imaging, we first confirmed that the photophysical behavior of purified FPs reproduces that observed in live cells, enabling in vitro investigation of photobleaching mechanisms. This validation allowed us to leverage in vitro spectroscopic analysis to identify distinct photoproducts contributing to fluorescence intensity loss—including non-absorbing species, species with new absorption bands, absorbing but non-fluorescent species, and fluorescent species with altered lifetimes—while avoiding the experimental limitations inherent to live-cell studies. Finally, structural characterization using SDS-PAGE and mass spectrometry provided unique chemical insights into the mechanisms driving the various photobleaching pathways.

## Methods

### Production and purification of fluorescent proteins

The production and purification of Histag-FPs, including Histag-Dronpa2, were carried out from *E. coli* bacteria using the pProEX-THa expression vector. The detailed protocol has already been described elsewhere^12^. For 500 mL of bacterial culture, approximately 10 mL of protein solution was obtained, at a concentration ranging between 100 µM and 400 µM. The proteins were stored in MCBtP buffer (2 mM CAPS, 2 mM MES, and 2 mM Bis–tris propane, pH 7.4) at −20°C. The sequences are available in SI, part A.

### Expression of fluorescent proteins in mammalian cells

Live-cell experiments were performed in COS7 cells. All FPs were cloned in pEGFP-N1 to be expressed as cytosolic proteins. Cells were cultivated in DMEM Glutamax (Invitrogen) supplemented with 5% FCS and antibiotics. For microscopy, cells were grown to 80% confluence on a glass coverslip (8 wells, Ibidi) 48 h before the experiment and transiently transfected 24 h prior to image acquisition with XtremeGene HP (Roche Diagnostics) following the supplier’s instructions. Cells were imaged in HBSS buffer. In Figures 2a and 2b, Aquamarine Field 1, mTurquoise, EYFP Field 1 and 2 were carried out at 26 °C using 10% of lamp input power for photobleaching (see below), while the other series were conducted at 20 °C using 20% of lamp input power.

### Preparation of FP-decorated Ni-NTA beads for microscopy experiments

To study the influence of the cellular environment on photobleaching, purified His-tagged proteins were immobilized onto Ni-NTA agarose beads with radii ranging from approximately 10 to 100 μm. A few microliters of FP stock solution were diluted into 1 mL of Ni-NTA bead suspension in a buffer containing 0.7 M NaCl, 30 mM NaH_2_PO_4_, 30 mM imidazole, pH 7.4 to obtain a final FP concentration of ~0.5 μM. The suspension was incubated for 30 min at 5 °C with constant stirring. Unbound FPs were then washed with Tris buffer (pH 8) by five centrifugation cycles (3 min at 5000 g). Beads were finally resuspended in 1 mL of the same Tris buffer (pH 8) and stored at 5 °C. For imaging experiments, a 50 μL droplet of the suspension was placed between two glass coverslips.

### Wide-field photobleaching and TCSPC-FLIM measurements

The microscope is equipped with a wide-field epifluorescence pathway on one side and TCSPC-based fluorescence lifetime imaging (FLIM) detection on the other, enabling successive measurements of fluorescence intensity and lifetime on the same field of view^13,14^. This setup was based on a TE2000 microscope with a 60×, 1.2 NA water-immersion objective (Nikon). The wide-field light source was employed to induce photobleaching of either purified Histag-FPs immobilized on Ni-NTA agarose beads or cytosolic FPs expressed in COS7 cells.

The wide-field pathway was equipped with a LED light source (SOLA, Lumencor) and an ORCA AG CCD camera (Hamamatsu). For each spectral selection, the irradiance was determined by actinometry (see SI, part B and Table S1). Photobleaching was triggered by continuous illumination and monitored by time-lapse image acquisition. The total experiment duration ranged from 1 to 10 min, depending on FP photostability. Image analysis was performed using ImageJ software. For each condition, 2–6 individual fluorescence intensity decay traces were collected, averaged, and fitted with a single-exponential function to extract the bleaching time constant (τ_bleach_) and the corresponding bleaching rate constant (k_bleach_).

The TCSPC pathway was equipped with 440 and 466 nm pulsed laser diodes (PicoQuant) driven by a PDL800 driver (20 MHz, PicoQuant) for CFPs, EGFP/YFPs, respectively. No pulsed excitation source was available for mCherry. Emitted fluorescence was spectrally filtered using a dedicated set of filters (see SI, part C and Table S4) positioned in front of an MCP-PMT detector (Hamamatsu Photonics). A C1 scanning head (Nikon) probed a 100 μm × 100 μm field of view. Signals were collected by a PicoHarp 300 TCSPC module (PicoQuant) and processed with SymPhoTime 64 software (PicoQuant). For each field of view, TCSPC fluorescence decays from all pixels above an intensity threshold were combined using the SymPhoTime 64 software to generate an averaged decay curve with reduced noise, representative of all cells in the field. For FLIM measurements of photobleached FPs, FLIM data were acquired throughout the photobleaching kinetics on the same field of view by alternating between the epifluorescence and TCSPC detection pathways of the setup (Figure S7).

### In vitro photobleaching and steady-state spectroscopy measurements

All absorption and fluorescence emission spectra, as well as photobleaching kinetics, were recorded using the custom-built setup, called BEAM, fully described and characterized previously^12^. Aquamarine, mTurquoise, and EGFP were bleached with a 445 nm laser diode (Oxxius LBX), whereas Citrine, EYFP, and mCherry were bleached with a 515 nm laser diode (Oxxius LBX). All photobleaching kinetics reported in this study have been conducted in 0.3×0.3 cm cuvette containing 35 µL of solution. To avoid the inner filter effect during fluorescence emission monitoring, the absorbance was kept below 0.1. At an absorbance of 0.5, the inner filter effect slightly alters the measured fluorescence intensity; therefore, the fluorescence intensity at the emission maximum was corrected using a custom workflow^12^ (Figure S9). The effective irradiances or photon fluxes corresponding to the irradiation conditions have been determined by actinometry (see SI, part B and Table S2). Irradiations were performed at 25 °C in Tris buffer (pH 8, chloride free).

### In vitro fluorescence lifetime measurements

Fluorescence decay curves were recorded using time-correlated single photon counting (TCSPC). The excitation source was a wavelength-tunable pulsed laser (Fianium) operating at a repetition rate of 10 MHz. The excitation wavelength (bandpass of 10 nm) was set to 435 nm for CFPs, 490 nm for EGFP, 515 nm for YFPs, and 585 nm for mCherry. Fluorescence emission was collected after a monochromator (bandwidth 6 nm) centered at 485 nm for CFPs, 510 nm for EGFP, 530 nm for YFPs, and 610 nm for mCherry. Fluorescence decay curves were detected using an MCP-PMT coupled to a PicoHarp 300 module (PicoQuant). The instrumental response function (IRF), obtained from scattered light of a LUDOX solution (DuPont), was typically 60–70 ps full width at half maximum (FWHM). Fluorescence decay curves and IRFs were recorded with approximately 5.10^6^ total counts per curve at typical acquisition rates of ~10^4^ counts per second. All samples were diluted to a maximum absorbance of 0.1 and thermostated at 20°C during acquisition. Data were collected using SymPhoTime software (PicoQuant).

### Fit of fluorescence decay curves and average lifetime

Fluorescence decay curves were analyzed using either SymPhoTime 64 software (PicoQuant) or Igor Pro software (Wavemetrics®). Unbleached samples were fitted with a mono-exponential function, while bleached samples were fitted with a bi-exponential function. For EGFP, which is known to intrinsically exhibit a biexponential decay with relatively close lifetimes (≈ 2 ns and 3 ns) corresponding to two configurations in equilibrium^15^, we first checked that this decay could be satisfactorily approximated by a single exponential, assuming that the two configurations contribute to a single population with an intermediate average lifetime (see Figures S8a and S8b). As no significant difference was observed using this approximation, we applied the same fitting procedure to EGFP as for the other FPs.

For in vitro experiments, one of the lifetimes was fixed at the value measured for the corresponding unbleached sample when fitting the decay of bleached samples:

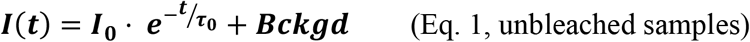

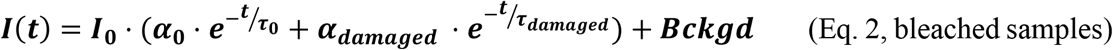

The average fluorescence lifetime < *τ* > is calculated as:

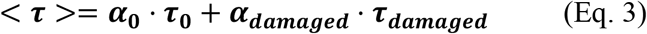

Where ***I***(***t***) is the transient fluorescence intensity, *Bckgd* the background fluorescence intensity, *I*_0_ a proportionality factor, ***α***_**0**_ and ***α***_***damaged***_ the normalized pre-exponential factors (***α***_**0**_ + ***α***_***damaged***_ = 1), ***τ***_**0**_ and ***τ***_***damaged***_ the lifetime of the unbleached and damaged species, respectively.

### SDS PAGE experiments

Sodium dodecyl sulfate polyacrylamide gel electrophoresis (SDS-PAGE) was performed using precast 10% acrylamide gels (NuPAGE 10% Bis-Tris Gel, Invitrogen), according to the manufacturer’s instructions and with the recommended reagents. Prior to loading, protein samples were denatured by mixing with the supplier’s sample buffer and heating at 90 °C for 10 min. For quantification, 8 µg of protein were loaded per well, which was low enough to avoid saturation of the main band intensity. For gels intended to visualize photoproduct bands—often much less intense than the main band—13 µg of protein was loaded per well. After electrophoresis, gels were colored with a solution containing Coomassie Brilliant Blue and stored in a solution containing 20% ethanol and 10% acetic acid before image acquisition for quantification using ImageJ software.

### Mass spectrometry experiments

Mass spectrometry (MS) experiments were performed on a QToF instrument (Synapt G2-Si, Waters Company, Manchester, UK). For direct infusion, samples were introduced in the electrospray ionization source (ESI) at a flow rate of 5 µL/min and analysed in positive ion mode over the mass-to-charge ratio (m/z) range of 500 to 5000 under denaturing conditions (30% acetonitrile/0.1% TFA). Instrumental and hardware parameters were optimized to maximize signal-to-noise ratio and were set as follows: capillary voltage = 3.5 kV, sampling cone = 80 V, source offset = 70 V, nebulization gas pressure = 5 bar, source temperature = 80 °C, desolvation temperature = 280 °C. Calibration was performed using sodium trifluoroacetate. The averaged RMS deviation was around 0.2 ppm. Mass predictions were performed using MassXpert2^16^. Raw data were extracted and pre-processed using MineXpert2^17,18^, mass deconvolution was performed with UniDec^19^ and deconvoluted spectra were analysed using Igor Pro (Wavemetrics®). For each sample, fractions of unmodified and oxidized FPs were determined from the relative intensity of each peak of the deconvoluted spectra after subtracting the spectra of the corresponding unbleached sample.

## Results

### Comparison of the variations in fluorescence intensity and fluorescence lifetime in living cells and *in vitro* during photobleaching

#### Actinometry and photobleaching indicators

Usual protocols for assessing FPs photostability typically monitor the decay kinetics of their fluorescence intensity under continuous irradiation, while maintaining constant experimental conditions, including excitation spectral range, irradiation power, illumination sequence, and the instrument used ^5,13,20^. Although the photostability hierarchy among FPs is generally maintained across setups, direct quantitative comparisons of photostability values remain challenging because irradiation conditions are often poorly characterized across them.

For instance, the photon flux *P*_*λ*_ (mol·m^−2^·s^−1^) received by the sample is not always precisely measured. In addition, excitation frequently occurs at a wavelength different from the protein’s absorption maximum. As a result, the actual quantity of absorbed photons per second, i_abs_ (mol·s^−1^), may be misestimated. To address this, we measured the photon flux at the experimental excitation wavelength, λ_ex_, 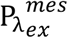, and corrected it to its equivalent at the absorption maximum, λ_max_, 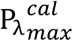, using the following equation:

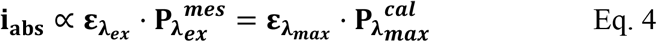

With 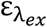 and 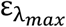, the molar extinction coefficients of the studied FP at *λ*_*ex*_ and *λ*_*max*_, respectively.

To enable a proper comparison between the wide-field microscopy and BEAM setups, the photon flux of each setup was carefully measured following an actinometric protocol inspired by Lahlou et al.^21^. In particular, we considered in the actinometry workflow that with BEAM, bleaching is triggered by lasers, which provide a monochromatic light compared to the filtered white light source used in the microscope (See SI, part B1). Our measurements show that the photon flux on the BEAM setup is 40 to 400 times lower than that of the microscope at the maximum power of its LED lamp (Tables S1 and S2).

In addition, it is crucial to have robust, standardized, and reproducible indicators for comparing the bleaching regimes in both setups. We previously developed a model linking the photobleaching rate constant, k_bleach_, to the photon flux, assuming that the photobleaching kinetics can be described by a monoexponential function ^12^ as confirmed experimentally (See SI, part B2). In this model, k_bleach_ is expressed as:

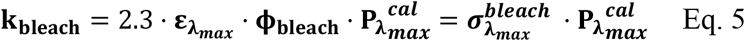

where 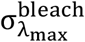 (m^2^·mol^−1^) is the photobleaching molar cross-section at *λ*_*max*_ and ϕ_bleach_, the photobleaching quantum yield.

The photobleaching molar cross-section, 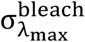, practically determines the photostability of FPs. Nevertheless, it depends on the FP’s ability to absorb photons efficiently through the molar extinction coefficient, 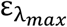 (Eq. 5). In imaging, FPs with high molar extinction coefficient are suitable to maximize brightness. Therefore, ideally, 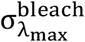 should be as low as possible and 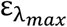 as high as possible, which can make interpreting the molar cross-section value somewhat counterintuitive. To provide an unambiguous comparison of FP photostability, we also computed the photobleaching quantum yield, ϕ_bleach_, where a low ϕ_bleach_ corresponds to high photostability.

#### Fluorescence intensity and photobleaching kinetics

The six FPs were investigated in live COS7 cells by wide-field microscopy and in vitro on BEAM for spectroscopic studies. In live-cell experiments, the photo-induced decrease in fluorescence intensity was monitored at five photon fluxes (Figure S3a). In vitro, photobleaching kinetics occurred on longer timescales than in live cells due to the lower photon flux and were therefore monitored at the maximum available photon flux (Figure S4). In all cases, the fluorescence intensity decreased monoexponentially with irradiation time, as previously reported for Citrine^12^. The parameters 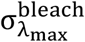 and ϕ_bleach_ were computed from the bleaching rate constant k_bleach_ (Figures S3, S4 and Table S3). From a quantitative point of view, only a limited number of studies have reported photobleaching quantum yield values for FPs and the available values show some variability^22– 24^. Nevertheless, the reported values consistently fall within the same order of magnitude, ranging from 10^−7^ to 10^−5^ depending on the FP studied, in excellent agreement with our measurements.

YFPs displayed the largest photobleaching molar cross-sections and were consequently the least photostable among the FPs examined (Figure 1a, Table S3). They were followed by the CFPs, and finally by EGFP and mCherry, which showed almost equivalent photostabilities. As the molar absorption coefficients of CFPs are significantly lower than those of the other FPs (See SI, part A), the comparison of photobleaching quantum yields favors the other FPs (Figure 1b). Nevertheless, YFPs remained the least photostable, consistent with prior reports^5,25^.

**Figure 1.**
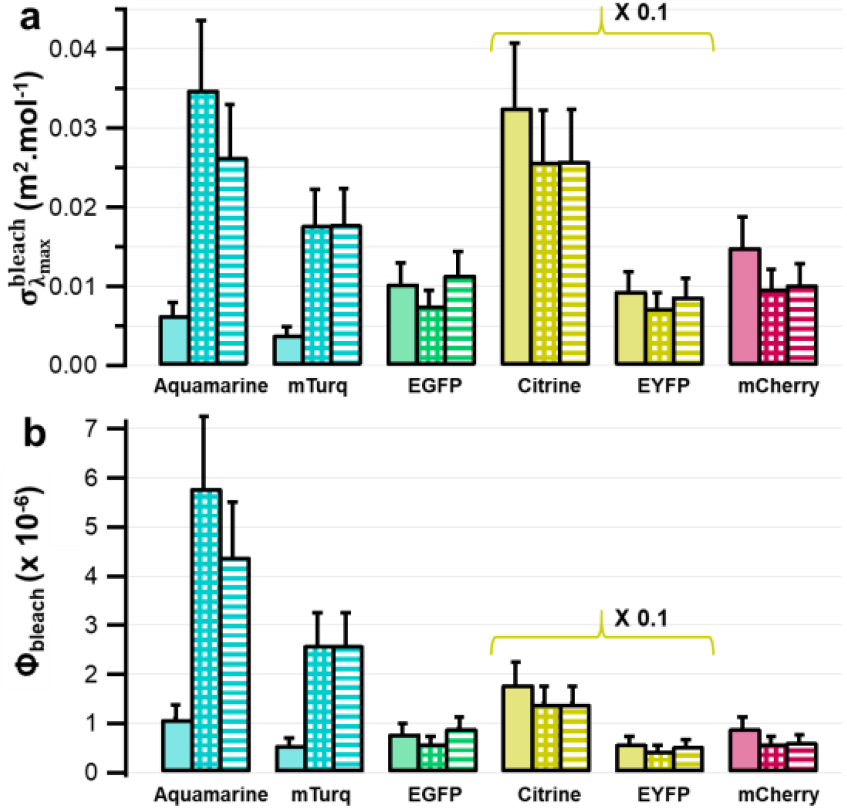
Photobleaching molar cross sections, 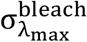 (m^2^·mol^−1^) (a) and photobleaching quantum yields, ϕ_bleach_ (b), for the six FPs investigated. Measurements were performed *in vitro* using BEAM (solid bars), and on a widefield microscope for purified FPs immobilized on Ni-NTA beads (crosshatched bars), or expressed in live COS7 cells (hatched bars). For clarity, values for YFPs (Citrine and EYFP) were divided by 10 in both panels. Error bars were estimated at 25% of the final values, primarily reflecting the relative uncertainty in photon flux measurements (20% as reported by Lahlou et al. ^21^), which constitutes the main source of error. An additional 5% was included to account for minor contributions from factors such as exponential fitting of photobleaching kinetics or spectral measurements of the microscope lamp.

Within each spectral family, substantial differences are observed despite minimal sequence variation. EYFP was found to be approximately three times more photostable than Citrine, and mTurquoise about 1.5 times more photostable than Aquamarine, despite differing by only a few amino acid substitutions — one residue between Citrine and EYFP, and four between Aquamarine and mTurquoise — highlighting the critical role of the chromophore environment in FP photostability.

The 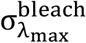 and ϕ_bleach_ values determined for Citrine, EYFP, EGFP, and mCherry are very similar in live cell under the microscope and *in vitro* with BEAM, despite differences in the photon flux between the two setups. In contrast, for CFPs, values are about 5 times lower *in vitro* (Figure 1). To assess whether this difference could arise from the cellular environment, the same parameters were measured for purified FPs immobilized on agarose Ni-NTA beads under the microscope. The resulting values closely match those obtained in live cells for all FPs, indicating that the cellular environment has no significant effect on the photobleaching kinetics (Figures 1, S3 and S5, Table S3).

Cranfill et al.^5^ proposed an empirical relationship linking the photobleaching rates to the excitation power of the light source I (in W), which is proportional to the photon flux:

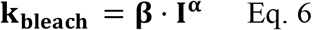

where **α** and **β** are constants. The coefficient α quantifies the deviation from linearity of the photobleaching rate constant with respect to excitation power. For YFPs, EGFP, and mCherry, α was reported to be close to 1, indicating an approximately linear dependence. This is consistent with our results, as both 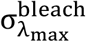 and ϕ_bleach_ were found to be independent of the excitation power for EGFP, YFPs, and mCherry over the power range explored. In contrast, for CFPs, a near-quadratic dependence of the photobleaching rate on excitation power has been reported^5^. Such nonlinearity directly affects the determination of 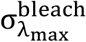 and ϕ_bleach_ when measurements are performed over large power range. In our experiments, photon fluxes ranged from 1 to 6 mol·m^−2^·s^−1^ under the microscope, whereas BEAM measurements were conducted at 0,013 mol·m^−2^·s^−1^, ie., approximately 75-450 times lower. Under these conditions, the differing 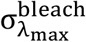 and ϕ_bleach_ values obtained for CFPs in the two setups likely arise from a shift in the photochemical reaction regime across the explored power range.

#### Fluorescence Lifetime variation during photobleaching

For all FPs, the effects of photobleaching on their fluorescence lifetime were evaluated in living COS7 cells using fluorescence lifetime imaging microscopy (FLIM) and compared with *in vitro* measurements performed with a TCSPC setup (Figures 2, and S6 to S8). Upon irradiation, the decay curves became more complex and the average fluorescence lifetimes decreased (Figure 2a,c, S6 and S8), indicating the formation of fluorescent photoproducts with shorter lifetimes. The average fluorescence lifetime decreased approximately linearly with fluorescence intensity loss, and FPs from the same spectral family followed comparable trends (Figure 2b, 2d). These behaviors were highly similar in live cells and *in vitro*, with the slope of lifetime loss relative to intensity loss conserved across both measurement conditions, consistent with previous reports^13,26,27^.

**Figure 2.**
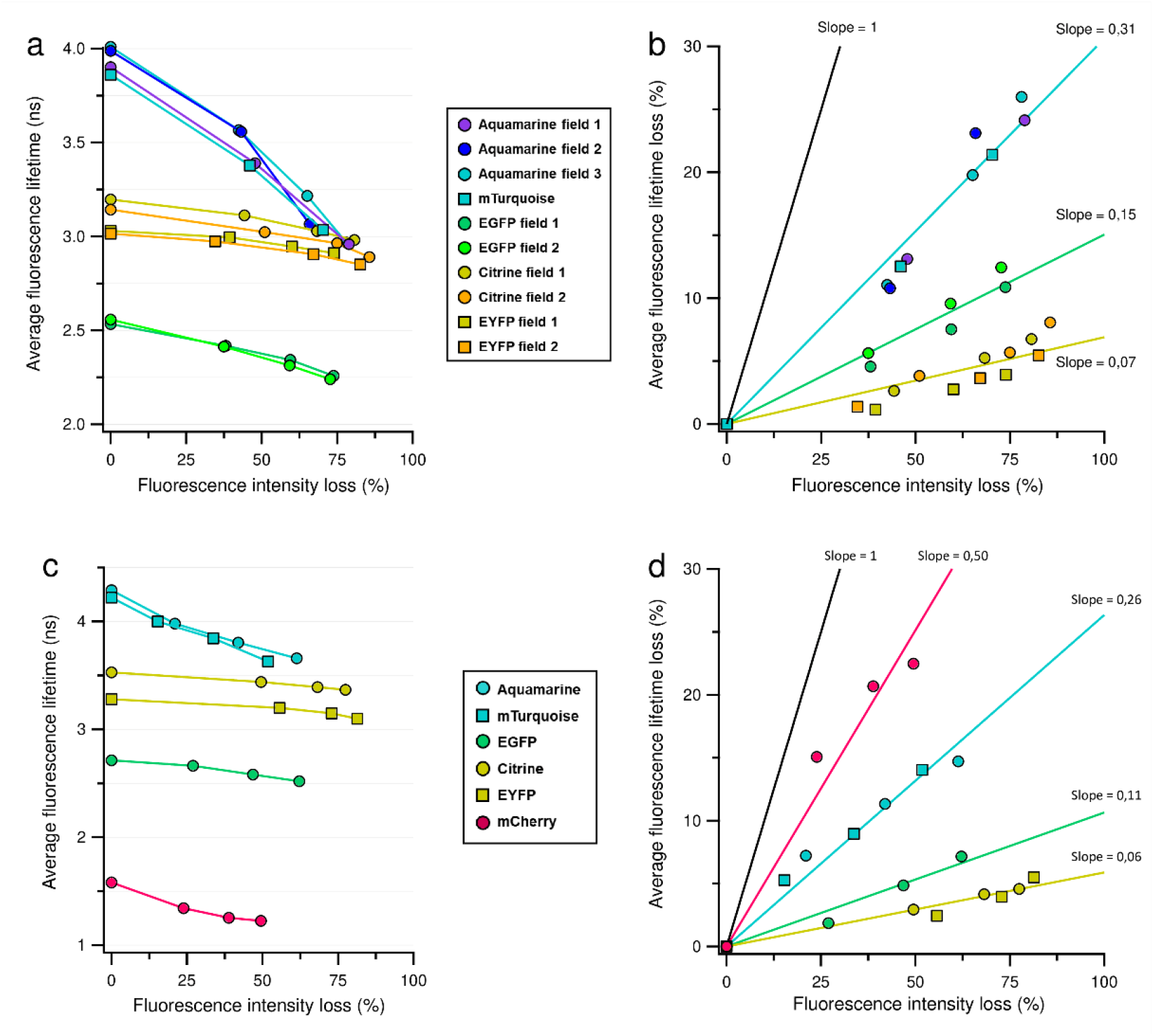
Average fluorescence lifetime of FPs as a function of the relative fluorescence intensity loss in live cells (a,b) and in vitro (c,d). Panels (a, c) show the average lifetime values. Panels (b, d) show the relative lifetime loss. Solid lines represent linear fits for each color family. The black line corresponding to the unit-slope line is shown as a visual guide. The FLIM setup is not equipped to excite mCherry.

### Spectroscopic study of FPs photobleaching in vitro

For most FPs, photobleaching indicators and fluorescence lifetime variations are consistent in live cells and *in vitro*, despite differences in excitation conditions. This supports the relevance of in vitro photobleaching experiments on purified FPs as representative of the photobleaching behavior observed in live cells. Such approaches enable physicochemical analyses that are experimentally inaccessible in living cells, thereby enabling a more detailed investigation of the molecular mechanisms governing photobleaching.

#### Evolution of Steady-State Absorption and Fluorescence Emission Properties During Photobleaching

A series of experiments was carried out at a concentration sufficiently high to record absorption and fluorescence emission spectra simultaneously during photobleaching (A_λex_=0.5, Figure 3, first and second columns). Across all the FP examined, a decrease in both maximum absorbance and fluorescence intensity was observed. However, the relative decay kinetics of absorption and emission differed depending on the FP (Figure 3, last column).

**Figure 3.**
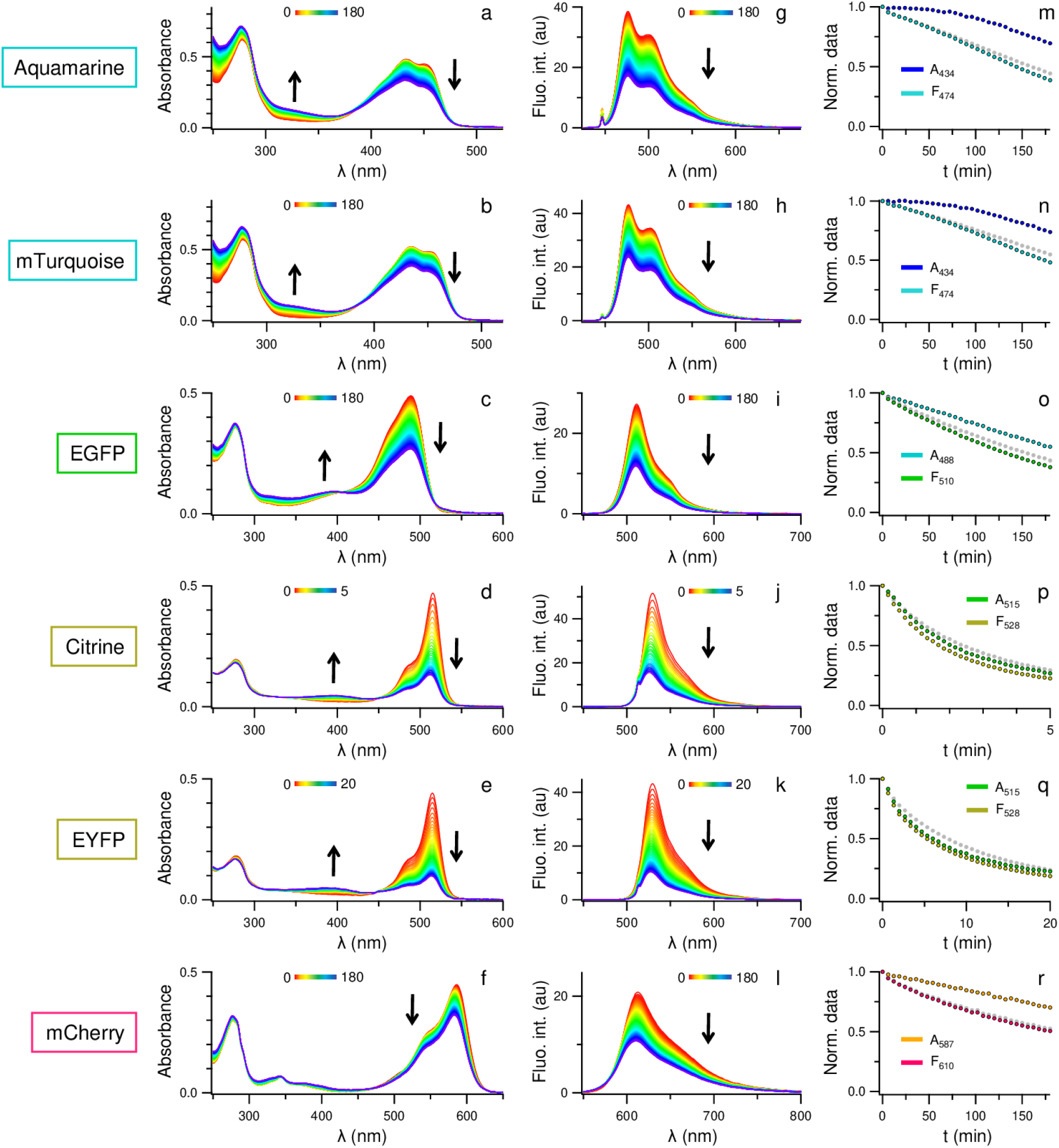
Time-dependent changes in absorption (left column) and fluorescence emission spectra (middle column), and corresponding photobleaching kinetics (right column). Each row corresponds to the results obtained for a given FP, indicated on the left. Irradiations were performed in vitro using lasers at 445 nm (irradiance: 0.39 W·cm^−2^) for CFPs and EGFP, and at 515 nm (irradiance: 0.60 W·cm^−2^) for YFPs and mCherry (Table S2). In the right column, gray dots represent the time-dependent fluorescence intensity at the wavelength of FP maximum emission during photobleaching, prior to inner filter effect correction (see Methods and Figure S9).

For YFPs, both the absorbance at 515 nm and the fluorescence intensity decreased with similar kinetics, indicating that photobleaching primarily results from a loss of the anionic chromophore’s ability to absorb, consistent with previous reports (Figures 3p,q)^9,12,28^. In contrast, CFPs showed only a slight decrease in absorbance compared to the marked drop in fluorescence intensity, suggesting that the chromophore remains largely chemically intact but partially loses its ability to fluoresce (Figures 3m,n).

EGFP and mCherry exhibited intermediate behavior: both the main absorption band and the fluorescence intensity decreased substantially, with fluorescence intensity decreasing more rapidly than absorbance (Figures 3o,r). These observations are in good agreement with the study by McLean et al. comparing ECFP, EGFP and EYFP^28^.

For all FPs, the absorbance and the position of the protein band at 280 nm remained constant, indicating that the environment of aromatic residues is mainly preserved during irradiation (Figures 2a-f)^29^. During the photobleaching of Citrine, EYFP, and EGFP, a new absorption band between 390 nm and 400 nm appears (Figures 2c-e). This band is consistent with the spectral signature of a protonated, non-fluorescent chromophore^30^ and suggests that certain photoproducts stabilize the protonated form of the chromophore, as previously observed in IrisFP^10^. However, considering the relative molar extinction coefficients of the protonated and anionic forms of these FPs^12,31,32^, protonation alone cannot account for the decrease in the anionic absorption band. In the case of CFPs, additional dark photoproducts were observed: a new absorption band appeared with a maximum between 320 nm and 340 nm (Figures 2a,b). To our knowledge, the corresponding chemical species has not been structurally characterized so far. Based on previous studies of tyrosine-derived chromophores exhibiting similar spectral signatures, this species could correspond to a hydrated form of the chromophore on the imidazolinone ring^33,34^. Precise quantification of this new band was limited by slight scattering in the UV region, consistent with the formation of larger assemblies, most likely multimers or aggregates.

To enable direct comparison of the shapes of the main absorption band and the fluorescence one, all spectra were normalized to unity at their respective maxima (Figure S10). For Citrine, EYFP and mCherry, a blue shift was observed in both absorption and emission bands. The contribution of the inner filter effect to these blue shifts was evaluated (Figure S11): these shifts are significant only for Citrine and mCherry and are likely due to subtle chemical modifications in the chromophore’s immediate environment. Interestingly, EYFP did not exhibit comparable photo-induced spectral shifts, despite differing from Citrine by only a single residue. In CFPs, a slight deformation of the absorption band was detected around 450 nm (Figures S10a,b). Under our irradiation conditions, no significant new emission bands, such as those associated with redding phenomena,^35^ were detected for any FP.

Altogether, these results highlight the diversity of reactive pathways and photoproducts leading to photobleaching, with their relative contributions varying across FPs. The identified photoproducts can be grouped into three main families (Scheme 2): (i) **dark** photoproducts, lacking both absorption and fluorescence; (ii) **colored** photoproducts, displaying new absorption bands but no detectable fluorescence; and (iii) photoproducts that retain normal absorption properties but exhibit altered or complex fluorescence emission characteristics.

Furthermore, photobleaching led to increasingly complex fluorescence decay kinetics, as well as reduced average fluorescence lifetimes (Figures 2, S6, and S8). The reduction in average fluorescence lifetime was relatively modest for YFPs and EGFP but markedly stronger for CFPs and mCherry (Figures 2a,c). This trend implies the additional formation of photoproducts that remain fluorescent yet display shorter fluorescence lifetimes, here referred to as **damaged** species. However, the formation of those species alone cannot explain the discrepancy between absorbance and fluorescence intensity decreases during photobleaching. We therefore introduced a last family of photoproducts in our model that retain normal absorption properties but exhibit negligible fluorescence, referred to as **dim** species (Scheme 2).

During photobleaching induced by white-light irradiation, the absorption spectra were recorded under steady-state conditions, while fluorescence emission was monitored using both time-resolved and steady-state methods (see SI part D). All measurements revealed similar qualitative trends, suggesting that the photochemistry described above is largely independent of the excitation wavelength.

**Scheme 2:**
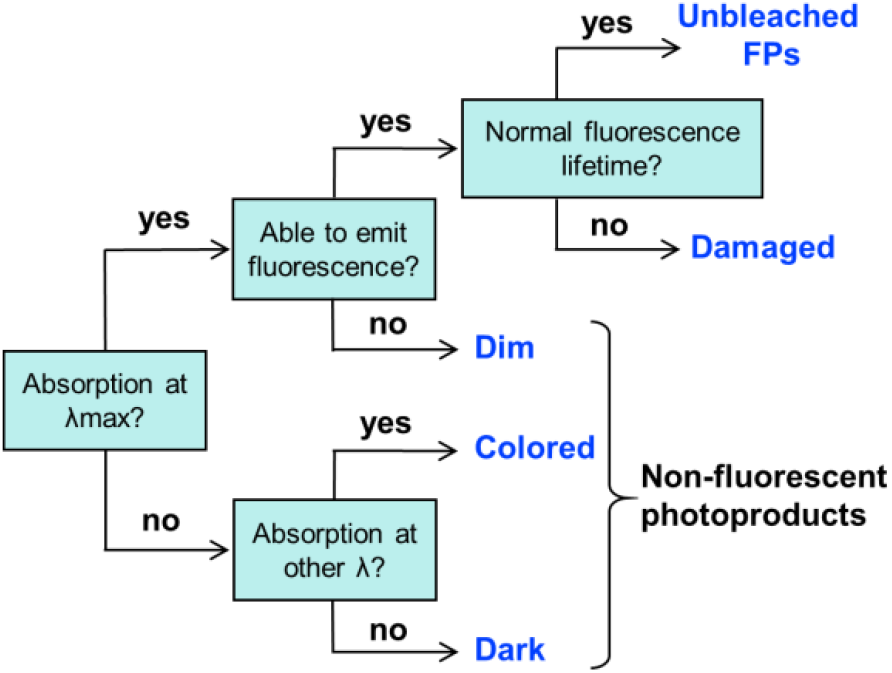
Classification of photoproduct populations formed in FPs upon photobleaching, based on their absorption characteristics and fluorescence emission abilities. Photoproducts are grouped into five classes: (i) **unbleached FPs** exhibiting normal absorption and fluorescence lifetime; (ii) **damaged species** retaining normal absorption but showing reduced fluorescence lifetime; (iii) **dim species** with normal absorption but negligible fluorescence emission; (iv) **colored photoproducts** showing new absorption bands without fluorescence; and (v) **dark photoproducts** lacking both absorption and fluorescence.

#### Quantification of the different photoproducts based on their spectroscopic signature

We developed a simple model to provide an estimate of the relative populations of photoproducts—unbleached, damaged, dim, colored, and dark—for the six FPs. This analysis used the complete set of spectroscopic data acquired during photobleaching, including absorbance, fluorescence intensity, and fluorescence lifetime measurements (see SI part E).

For example, to quantify the fraction of damaged proteins within the fluorescent FP population, fluorescence decay curves were fitted using biexponential functions, fixing the longer lifetime component to the FP lifetime prior to photobleaching. The relative amplitudes of the longest and shortest lifetime components corresponded directly to the proportions of unbleached and damaged species, respectively (Table S5).

Photoproducts populations were calculated at increasing photobleaching levels, highlighting distinctive patterns across FP families (Figure 4). YFPs predominantly formed dark and colored photoproducts, with minimal changes across bleaching levels. In contrast, CFPs exhibited a more balanced distribution among all photoproduct categories, with a significant proportion of damaged molecules at low photobleaching levels that progressively converted into dark and colored species as the bleaching level increased. mCherry behaved similarly to CFPs, but no dim species were detected. EGFP resembled YFPs but showed a small fraction of damaged species. In addition, Citrine photobleaching generated roughly equal amounts of dark and colored photoproducts, whereas EYFP produced about 20% more colored species, suggesting a higher propensity for chromophore protonation in EYFP during photobleaching.

**Figure 4.**
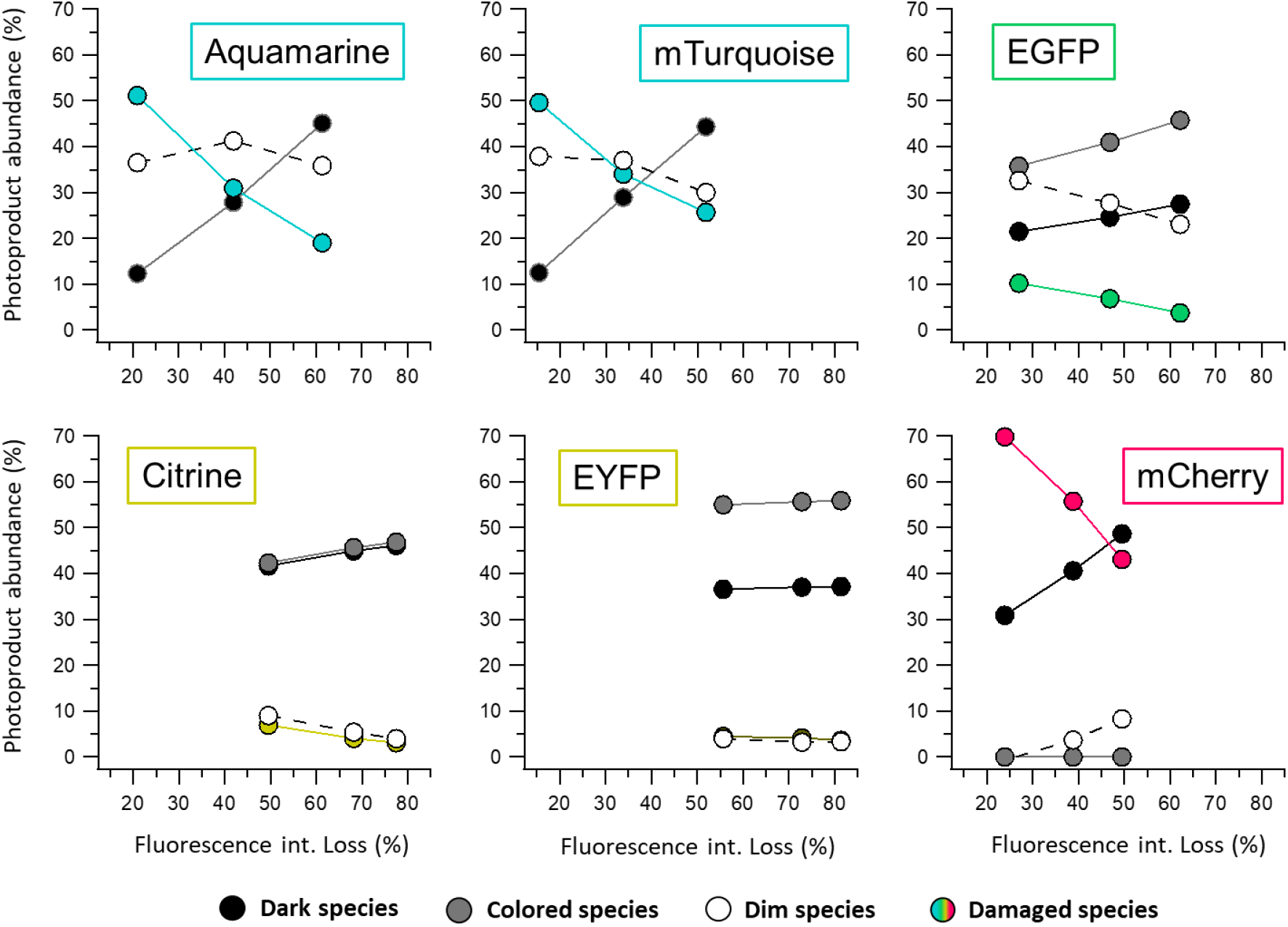
Fractions of photoproduct abundances in FP solutions at varying photobleaching levels. Fractions represent the amount of each species relative to the total photoproduct abundance. For CFPs, the dark and colored populations were pooled together (dark dots with grey stroke), as the photoproduct with a main absorption band at 340 nm could not be quantified individually.

### Description of FPs photobleaching at molecular scale

The previous experiments enabled assignment photophysical properties to the different photoproducts formed during photobleaching and quantification of their relative proportions. To gain molecular-level insight into the underlying chemical processes, bleached samples were analysed by SDS-PAGE and mass spectrometry. Furthermore, we calculated the cumulative photon dose delivered to each FP sample as the product of the equivalent photon flux 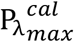 and the irradiation time. This parameter corrects for differences in photon flux received by FPs of distinct spectral families, enabling direct comparison across them. It therefore reflects not only the propensity of each FP to generate specific photoproducts, such as cleavage fragments or dimers, but also its ability to form these photoproducts rapidly under a given photon dose.

#### Three families of chemical modifications identified by SDS-PAGE

For all FPs studied, SDS-PAGE gels showed a broadening of the main band around ~30 kDa, indicating molecular heterogeneity likely arising from the formation of photoproducts with molecular weights close to that of the parent FP. Two additional lower-molecular-weight bands, consistent with photocleavage products, and a band at ~60 kDa, likely corresponding to covalent dimers, were also observed (Figures 5a and S17). For mCherry, a substantial fraction (~25%) of cleaved proteins was detected independently of photobleaching. This behavior originated from an autolysis reaction induced by heating prior to SDS-PAGE, as previously described for DsRed, the ancestor of mCherry^36^.

**Figure 5.**
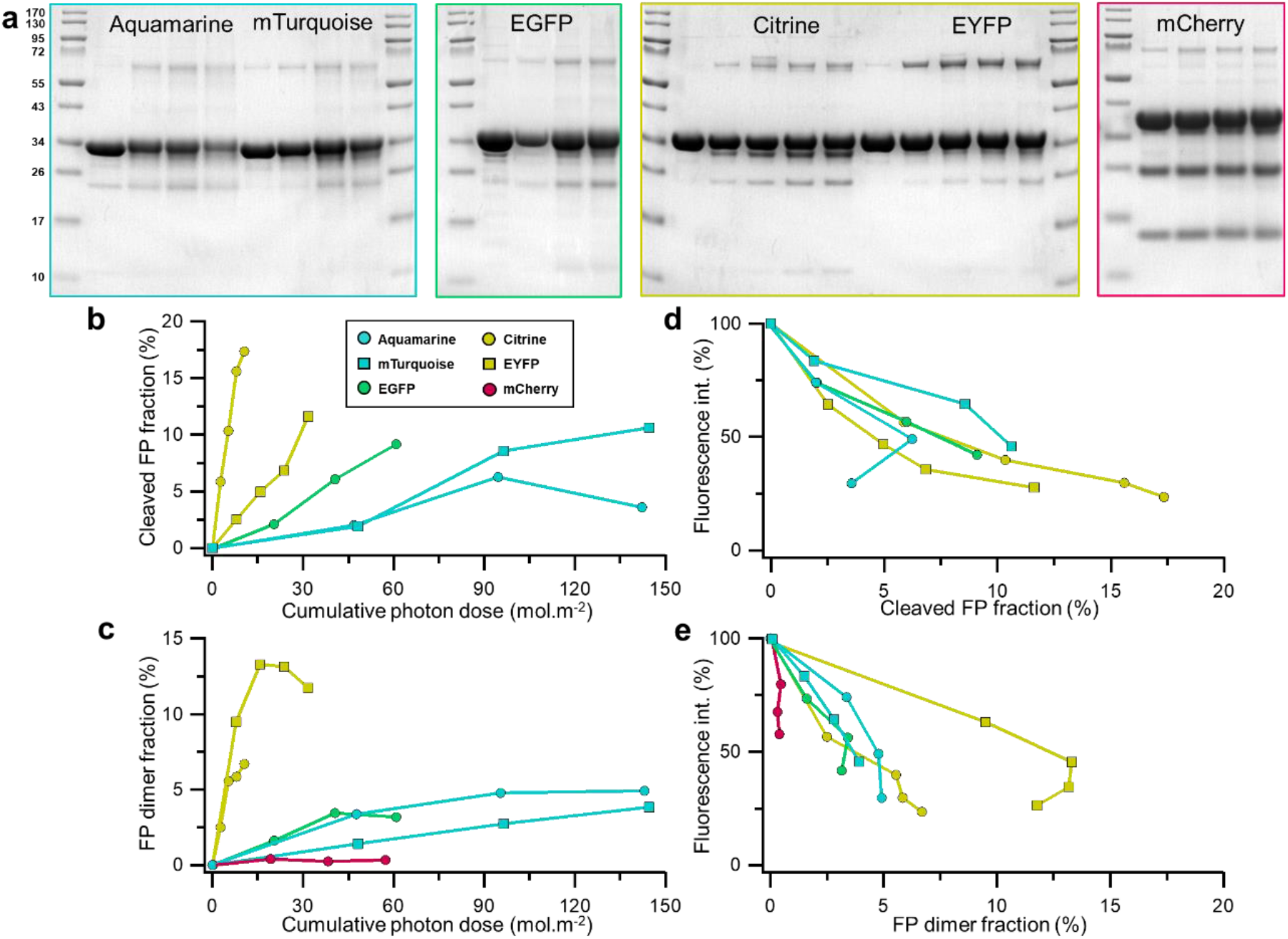
Molecular consequences of photobleaching. a) SDS-PAGE of irradiated FP samples. Molecular mass markers of the protein ladder are indicated in kDa. The fractions of cleaved FP and FP dimer as a function of cumulative photon dose are shown in b) and d), respectively. The dependence of fluorescence intensity on the fractions of cleaved FP and FP dimer is shown in c) and e), respectively. For clarity, the fractions of cleaved and dimeric FPs present prior to irradiation were systematically subtracted from the values measured during irradiation.

The evolution of the cleaved fraction depended strongly on the cumulative photon dose (Figure 5b). At equivalent doses, Citrine underwent more extensive cleavage than EYFP, EGFP, and CFPs. Across the FPs examined here, the fraction of photocleaved FPs correlated with the fluorescence intensity loss (Figure 5c), suggesting that cleavage contributes to bleaching. Nevertheless, cleavage never accounted for more than ~25% of total photoproducts. The highest level was observed for Citrine, with 17% of cleaved proteins at 77% total fluorescence intensity loss. Even assuming all cleaved species were non-fluorescent, photocleavage would therefore explain only ~ 22% of the overall photobleached species. With the exception of mCherry, all FPs studied displayed a clear tendency to form dimers upon photobleaching (Figure 5d). Nevertheless, dimerization remained well below 10% in most cases, except for EYFP, which reached nearly 15% and significantly impacted its photostability (Figure 5e).

#### Detailed analysis of the FP modifications by mass spectrometry

Mass spectrometry analysis of the bleached sample under denaturing conditions confirmed the SDS-PAGE observations (Figure S18). Pronounced molecular heterogeneity was observed around 30 kDa (Figures 6 and S19) together with covalent dimers (Figure S20). The data confirmed qualitatively that EYFP was more prone to dimerization than the other FPs examined (Figure S20). In addition, both the intensity and multiplicity of dimer mass signals increased progressively with photobleaching level, consistent with the formation of multiple distinct dimeric species during photobleaching (Figure S20).

**Figure 6.**
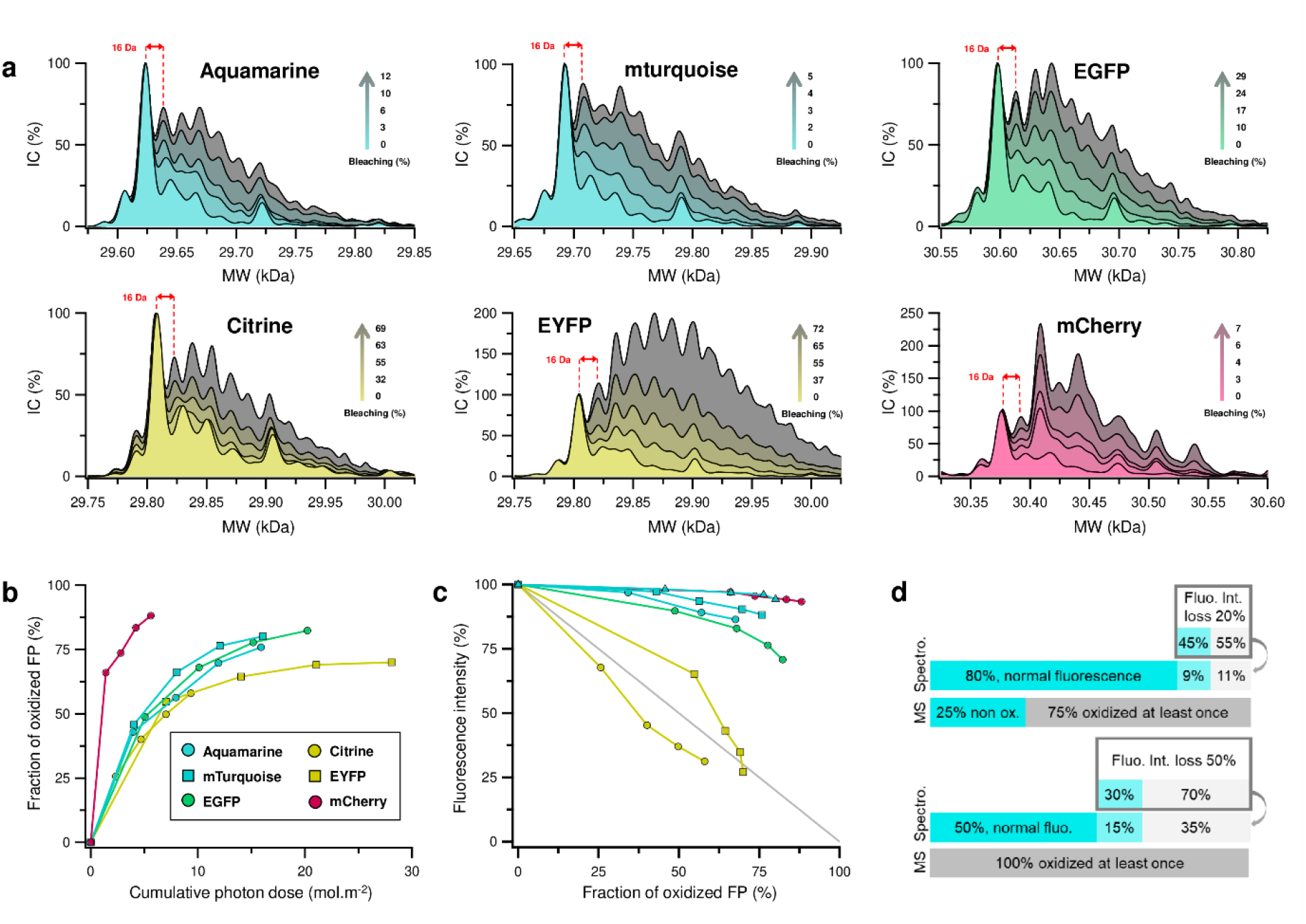
Mass spectrometry analysis of photobleached purified FPs in solution. a) Deconvoluted mass spectra of FPs recorded at increasing photobleaching level. Each spectrum is normalized to 100 % for the unmodified FP peak. b) Fraction of FPs oxidized at least once as a function of the cumulative photon dose (mol·m^−2^), determined from the relative peak intensities in panel (a). c) Fluorescence intensity as a function of the fraction of oxidized FPs at least once. d) Comparison of the proportions of fluorescent (blue) and non-fluorescent (grey) mTurquoise derived from the spectroscopic analysis (Figure 4) with the proportion of molecules oxidized at least once, as determined by the mass spectrometry analysis (panel c). This comparison is presented for fluorescence intensity losses of 20% and 50%.

The molecular heterogeneity at masses corresponding to full-length FPs was particularly pronounced for CFPs and mCherry, and to a lesser extent for EGFP. The signal corresponding to the unmodified FPs nearly vanished once photobleaching exceeded 30% (Figure S19). In contrast, for Citrine and EYFP, unmodified FPs remained detectable even above 60% photobleaching (Figure 6a).

At bleaching levels where photoproducts around 30 kDa were observed, successive mass increments of +16 Da appeared, consistent with multiple photo-oxidation events (Figure 6a). Oxidation signatures differed between FPs: mCherry exhibited multiple mass increments, predominantly of +32 Da, suggesting distinct photochemical pathways, whereas Citrine showed –16 Da mass shifts, likely arising from specific oxidation reactions^37^.

Regardless of their intrinsic photostability, all FPs underwent extensive oxidation upon irradiation. The fraction of FPs oxidized at least once consistently reached a plateau above 60% (Figure 6b). Among the proteins studied, mCherry generated the largest number of oxidized species (Figure 6b). In all cases, this oxidized fraction exceeded that of cleaved or dimerized FPs, highlighting oxidation as the dominant light-induced chemical pathway. The evolution of the average number of oxidation events paralleled the increase in the fraction of FPs oxidized at least once (Figures 6b and S21), suggesting statistically distributed oxidation events.

In addition, the relationship between oxidation and fluorescence intensity loss varied considerably among FPs (Figure 6c). As a general trend, the more photostable FPs —CFPs, and mCherry—required a high fraction of oxidized species before substantial fluorescence intensity loss occurred, implying that many oxidized forms can remain fluorescent. mTurquoise provides a clear illustration of this behavior. At 20% photobleaching, approximately 75% of the FPs have already undergone at least one oxidation event, whereas only 20 % of the population exhibits altered fluorescence properties (Figure 4). Among these photoproducts, roughly half correspond to fluorescent species with shortened lifetimes (damaged species), while the other half are dim or dark. Thus, oxidation can lead either to damaged fluorescent species or to non-fluorescent species, while a large fraction of oxidized molecules remains photophysically intact. At higher bleaching levels (≈50%), fluorescent damaged species account for about 30% of the photoproducts, whereas the remaining bleached population consists predominantly of dark or dim species. At this stage, the proteins are already extensively oxidized (Figure S19) with only minor contributions from backbone cleavage (7%) or dimerization. Overall, these observations indicate that oxidation populates both lifetime-shortened fluorescent species and predominantly non-fluorescent species.

In contrast, for YFPs, fluorescence intensity loss was nearly proportional to oxidation: ~50% of oxidized FPs correspond to ~50% non-fluorescent FPs, indicating that species oxidized at least once are largely non-fluorescent (Figure 6c). EGFP, once again, displays an intermediate behavior in both its fluorescence and oxidation response.

## Discussion

### Different mechanisms and the influence of the FP

#### Chromophore- and FP-dependent photobleaching

The photochemical behavior of FPs is governed by both the nature of the chromophore and the structural features of the protein scaffold. In CFPs, the chromophore is tryptophan-based, whereas in YFPs, EGFP and mCherry, it is tyrosine-based. In addition, the sequence —and consequently the β-barrel scaffold— of mCherry, derived from a different marine organism, differs substantially from that of the other FPs examined here (see SI part A). In our experiments, fluorescence intensity loss proceeds through three common chemical mechanisms—oxidation, dimerization, and peptide-backbone cleavage. Their relative contributions, however, vary substantially among FPs, leading to distinct behaviors under irradiation, including variations in photobleaching rates, photoproduct formation, and spectroscopic changes. In CFPs, photobleaching primarily reflects a partial loss of quantum yield (damaged species), that evolves into dark and dim species at high bleaching levels. In contrast, in YFPs, photobleaching is accompanied by a rapid loss of absorption at 515 nm, concomitant with the decrease in fluorescence intensity, suggesting that early photochemical modifications disrupt the chromophore’s π-conjugated system.

#### Chemical pathways of oxidative photobleaching

The most common mechanism of photobleaching in organic chromophores involves photosensitization by molecular oxygen, which can generate reactive oxygen species (ROS), including singlet oxygen, through diverse pathways^38^. The involvement of molecular oxygen in FP photobleaching has been demonstrated in GFP^39–41^ and in the red FP KillerRed^42–44^ through the detection of oxygen-derived photoproducts. Access of oxygen to the chromophore cavity is therefore required; Although this access is sterically limited by the β-barrel, it is also necessary for chromophore maturation.

In our experiments, multiple oxidation events detected by mass spectrometry reveal that oxidative processes affect not only the chromophore but also the protein scaffold. The spectroscopic changes in absorption, fluorescence emission, and lifetime of Aquamarine and mTurquoise, as well as the modification of their mass spectra, closely resemble those we previously observed for ECFP directly exposed to HO^•^ generated by gamma radiolysis^45,46^. The observed chemical modifications are consistent with reactions expected under oxidative stress conditions and are well documented for free amino acids, peptides, and proteins, particularly those containing methionine or aromatic residues^37,47,48^. The sensitivity of FPs fluorescence to H_2_O_2_ has also been reported for many fluorescent proteins, particularly EGFP and EYFP^49^.

#### Structural and photophysical consequences of oxidation

Oxidative modifications in FPs can perturb the structural integrity of the β-barrel, altering its rigidity and increasing the flexibility of the chromophore’s immediate environment. These changes introduce additional degrees of freedom around the chromophore, favoring non-radiative de-excitation pathways, and can shorten the fluorescence lifetime ^50,51^. Consistent with this mechanism, the damaged species in our experiments exhibited substantially shorter lifetimes, ranging from 30 to 64% of the native FP’s fluorescence lifetime. This suggests a partial correspondence between oxidized FPs and lifetime-shortened species, although oxidation does not necessarily lead to immediate fluorescence intensity loss. A striking example is mCherry, which is highly prone to oxidative modifications. In this protein, damaged species exhibit particularly short fluorescence lifetimes, yet extensive oxidation results in only a modest decrease in fluorescence intensity under our irradiation conditions. This observation underscores that chemical stability is not directly correlated with fluorescence intensity. Such behavior is consistent with the proposed biological role of FPs in corals, where they may act as photoprotective pigments that buffer ROS generated in their environment^52^.

#### Photoinduced dimerization of fluorescent proteins

Proteins exposed to oxidative stress are known to dimerize through a variety of oxidation-driven mechanisms, most commonly via covalent cross-links between tyrosine or tryptophan residues^53,54^. Although the mass resolution of our spectra was insufficient to assign precise molecular weights, the observed mass shifts suggest that the detected dimers are oxidized. To our knowledge, photoinduced dimerization of FPs has not previously been reported, and in our case, it appeared as a minor byproduct, most likely arising from oxidation. This interpretation is consistent with the relatively slow accumulation of dimers observed during irradiation. The ROS responsible for oxidation are generated near the chromophore and must first diffuse through the protein matrix toward the solvent-exposed surface before reaching residues capable of forming intermolecular cross-links with neighboring FPs. Empirically, dimerization also appears to be concentration-dependent in YFPs, which is consistent with a mechanism involving intermolecular oxidative coupling.

#### Photochemical peptide-backbone cleavage

FPs appear to contain a reactive site at the chromophore that can undergo light-induced cleavage of the polypeptide chain. Such a cleavage is often associated with photoconversion of proteins, including Kaede^55,56^, EosFP^57,58^, and IrisFP^59^. In these proteins, a photochemical reaction induces backbone cleavage near the chromophore, extending its π-conjugated system and resulting in a color change. However, this type of reaction has rarely been associated with photobleaching and has been reported once in EGFP^8^. In this study, depending on the protein, cleavage just upstream of the chromophore likely proceeds via a photochemical pathway and results in complete fluorescence intensity loss, thereby contributing to the formation of dark species.

#### Influence of sequence and key residues on photobleaching

Our study highlights that mutations in the protein sequence affected both the nature of photoproducts and the rates of photobleaching. Significant differences were observed within CFPs and YFPs themselves, underlining the critical role of specific amino acids in modulating the chromophore’s reactivity toward light. A striking example is the comparison between EYFP and Citrine. EYFP exhibited higher photostability than Citrine, with a photobleaching molar cross-section and quantum yield roughly 2–3 times lower. These proteins differ by only a single mutation (Q69M), the reduced photostability of Citrine may be related to the reactivity of methionine with ROS^37,45,47,48,60^. Supporting the role of oxidation in photobleaching, X-ray diffraction analysis of a photobleached IrisFP crystal revealed photoinduced oxidation of a methionine residue, which stabilized a protonated, non-fluorescent form of the chromophore^10^. Nevertheless, despite its low photostability, Citrine is more widely used and better suited for live-cell imaging: it is 30% brighter than EYFP^3^ and its fluorescence properties are less sensitive to pH and chloride ion concentration^20^. Another example is the comparison between mTurquoise and Aquamarine. These two CFPs differ by only four residues at positions 72, 148, 175, and 206 (Figure 1), yet mTurquoise exhibited greater photostability. Interestingly, Aquamarine does not contain residues particularly prone to oxidation compared to mTurquoise. The observed differences in photostability may therefore arise primarily from the structural consequences of oxidation, which can increase the flexibility of the chromophore’s immediate environment, thereby enhancing non-radiative de-excitation pathways that compete with fluorescence.

#### Summary and Implications of Photobleaching Mechanisms

Taken together, these results highlight the diversity of photobleaching chemical mechanisms across FPs and their distinct spectroscopic consequences. Although many of these mechanisms are common to different FPs, their relative contributions vary considerably and appear to depend primarily on FP color, consequence of the chemical structure of the chromophore and its immediate environment. By contrast, photobleaching kinetics seem to be more strongly influenced by the overall protein sequence, and can differ significantly even among FPs of the same color. Beyond the fundamental interest of distinguishing these mechanisms, this divergence is of practical importance, as it gives rise to distinct photophysical signatures, including changes in fluorescence intensity, spectrum, and lifetime, which can ultimately lead to different experimental outcomes in fluorescence microscopy.

### Consequences of FPs photobleaching for imaging

#### Impact of Photobleaching on Fluorescence Imaging

One of the key challenges in elucidating the mechanisms and structural determinants of FP photobleaching lies in its impact on imaging performance. During fluorescence imaging experiments aimed at visualizing cellular compartments or tracking proteins over time, photobleaching progressively reduces fluorescence intensity, thereby limiting the duration of observation and the achievable temporal resolution. In addition, the decrease in emitted signal lowers the signal-to-noise ratio which can degrade image quality and reduce effective spatial resolution. Photobleaching is strongly dependent on the illumination conditions characteristic of each fluorescence microscopy modality. For example, the much higher average irradiances typically used in confocal microscopy (> 1 kW·cm^−2^) compared with wide-field imaging (< 1 W.cm^−2^) lead to faster photobleaching^61^. Techniques enabling imaging beyond the diffraction limit, such as Stimulated Emission Depletion (STED) microscopy, require even higher irradiances^62^. These conditions can be particularly damaging for FPs, even though the depletion beam is not delivered at the chromophore absorption maximum required to generate the STED effect^63^. Beyond the simple loss of fluorescence signal, photobleaching can also generate reactive oxygen species (ROS), which can diffuse and damage nearby proteins, including those not directly illuminated. This phenomenon is exploited in Chromophore-Assisted Light Inactivation (CALI) to generate localized high ROS concentrations^64^.

#### Impact of photobleaching on quantitative FLIM FRET imaging

For quantitative imaging, the impact of photobleaching cannot be overlooked. Photobleaching directly reduces fluorescence intensity, limiting the accuracy of intensity-based imaging methods. Our results further show that fluorescence lifetimes, monitored by FLIM, are also influenced by photobleaching, consistent with previous observations on ECFP, EGP, EYFP and mCherry^27^. When FLIM is used to determine the Förster Resonance Energy Transfer (FRET) efficiency between donor and acceptor fluorophores, photobleaching can lead to misinterpretation. FRET efficiency depends on spectral overlap between donor emission and acceptor absorption, as well as the distance and relative orientation of the fluorophores. Any variation in donor/acceptor geometry (distance or orientation) affects the FRET signal, which is exploited to monitor protein–protein interactions or the response of FRET-based biosensors. FRET reduces the donor lifetime by introducing an additional non-radiative de-excitation pathway from the excited state. In cyan/yellow FRET pairs, donor photobleaching (CFP) reduces the fluorescence lifetime independently of any FRET modulation, leading to an overestimation of FRET efficiency. On the acceptor side (YFP), photobleaching decreases the main absorption band, thereby reducing spectral overlap with the donor emission and, consequently, reducing FRET efficiency, regardless of any geometrical change. According to our results, similar artifacts may also occur when using EGFP as the donor and mCherry as the acceptor, although the impact should be smaller.

## Conclusion

This work proposes a framework to quantitatively study FP photobleaching. The analytical workflow combines mass spectrometry, spectroscopic, and biochemical approaches to characterize photoinduced chemical modifications in FPs during photobleaching and their consequences for photophysical properties and imaging performance. We demonstrate that these modifications include a combination of oxidation, dimerization, and, peptide backbone cleavage upstream of the chromophore. The extent and nature of these modifications and their photophysical consequences strongly depend on the FP sequence, the chromophore structure, and its surrounding protein environment. These aspects were examined in detail across six FPs derived from two marine organisms, bearing different chromophore types, and covering four emission wavelength ranges. The range of photon flux densities used in this study corresponds to irradiation conditions encountered in wide-field microscopy experiments.

More specifically, we reveal a molecular diversity of photoproducts with distinct photophysical properties that can still contribute to fluorescence and/or absorption signals. As a result, for imaging experiments, photobleaching in fluorescent proteins is more complex than a simple ON–OFF transition where the signal is completely extinguished. Furthermore, in addition to fluorescence intensity loss, we quantified the photobleaching-induced decrease in fluorescence lifetimes. These photophysical changes are particularly relevant for FLIM and FRET experiments, as they can lead to misinterpretations and introduce quantitative bias into experimental readouts.

This study also emphasizes that the concept of photostability is strongly context-dependent and highlights the difficulty of comparing the photostability of different FPs under irradiation conditions involving different photon fluxes and spectral settings. To address this, we defined two indicators 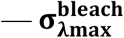 and **ϕ**_**bleach**_ — that enable accurate, quantitative comparisons of FP performance across different instruments.

Taken together, these results provide new insight into the molecular mechanisms of FP photobleaching and their consequences for quantitative fluorescence imaging.

## Acknowledgments

This work was also supported by LabEx PALM, Grant ANR-10-LABX-0039-PALM and BioProbe-France 2030 program Grant ANR-11-IDEX-0003. We thank A. Lahlou and T. Le Saux for their advice and the gift of the plasmid coding for Dronpa2.

