## Supplementary Information for "« Fluorescent Protein Photobleaching: From molecular processes to spectromicroscopy »"

### Supporting Information
### Fluorescent Protein Photobleaching:
### From molecular processes to spectromicroscopy

*Théo Beguin, Kaixin Wang, Yasmina Bousmah, Ninette Abou Mrad, Frederic Halgand, Hélène Pasquier, Marie Erard**

Université Paris-Saclay, CNRS, Institut de Chimie Physique, UMR CNRS 8000, 91405, Orsay, France.

#### A) Protein sequences and general characteristics

**Aquamarine**

**MW = 29625 Da**
**$\varepsilon_{430nm}$ = 26000 $M^{-1}\cdot cm^{-1}$**

(Histag) MSYYHHHHHHDYDIPTTENLYFQGA
(1) VSKGEELFTGVVPILVELDGDVNGH
(26) KFSVSGEGEGDATYGKLTLKFICTT
(51) GKLPVPWPTLVTTLSWGVQCFSRYP
(76) DHMKQHDFFKSAMPEGYVQERTIFF
(101) KDDGNYKTRAEVKFEGDTLVNRIEL
(126) KGIDFKEDGNILGHKLEYNYISGNV
(151) YITADKQKNGIKANFKIRHNIEDGS
(176) VQLADHYQQNTPIGDGPVLLPDNHY
(201) LSTQSALSKDPNEKRDHMVLLEFVT
(226) AAGITLGMDELYK

**mTurquoise**

**MW = 29694 Da**
**$\varepsilon_{434nm}$ = 30000 $M^{-1}\cdot cm^{-1}$**

(Histag) MSYYHHHHHHDYDIPTTENLYFQGA
(1) VSKGEELFTGVVPILVELDGDVNGH
(26) KFSVSGEGEGDATYGKLTLKFICTT
(51) GKLPVPWPTLVTTLSWGVQCFARYP
(76) DHMKQHDFFKSAMPEGYVQERTIFF
(101) KDDGNYKTRAEVKFEGDTLVNRIEL
(126) KGIDFKEDGNILGHKLEYNYISDNV
(151) YITADKQKNGIKANFKIRHNIEDGG
(176) VQLADHYQQNTPIGDGPVLLPDNHY
(201) LSTQSKLSKDPNEKRDHMVLLEFVT
(226) AAGITLGMDELYK

**EGFP**

**MW = 30598 Da**
**$\varepsilon_{488nm}$ = 56000 $M^{-1}\cdot cm^{-1}$**

(Histag) MSYYHHHHHHDYDIPTTENLYFQGA
(Histag) MDPPVATM
(1) VSKGEELFTGVVPILVELDGDVNGH
(26) KFSVSGEGEGDATYGKLTLKFICTT
(51) GKLPVPWPTLVTTLTYGVQCFSRYP
(76) DHMKQHDFFKSAMPEGYVQERTIFF
(101) KDDGNYKTRAEVKFEGDTLVNRIEL
(126) KGIDFKEDGNILGHKLEYNYNSHNV
(151) YIMADKQKNGIKVNFKIRHNIEDGS
(176) VQLADHYQQNTPIGDGPVLLPDNHY
(201) LSTQSALSKDPNEKRDHMVLLEFVT
(226) AAGITLGMDELYK

**Citrine**

**MW = 29808 Da**
**$\varepsilon_{515nm}$ = 77000 $M^{-1}\cdot cm^{-1}$**

(Histag) MSYYHHHHHHDYDIPTTENLYFQGA
(1) VSKGEELFTGVVPILVELDGDVNGH
(26) KFSVSGEGEGDATYGKLTLKFICTT
(51) GKLPVPWPTLVTTFGYGLMCFARYP
(76) DHMKQHDFFKSAMPEGYVQERTIFF
(101) KDDGNYKTRAEVKFEGDTLVNRIEL
(126) KGIDFKEDGNILGHKLEYNYNSHNV
(151) YIMADKQKNGIKVNFKIRHNIEDGS
(176) VQLADHYQQNTPIGDGPVLLPDNHY
(201) LSYQSALSKDPNEKRDHMVLLEFVT
(226) AAGITLGMDELYK

**EYFP**

**MW = 29805 Da**
**$\varepsilon_{515nm}$ = 67000 $M^{-1}\cdot cm^{-1}$**

(Histag) MSYYHHHHHHDYDIPTTENLYFQGA
(1) VSKGEELFTGVVPILVELDGDVNGH
(26) KFSVSGEGEGDATYGKLTLKFICTT
(51) GKLPVPWPTLVTTFGYGLQCFARYP
(76) DHMKQHDFFKSAMPEGYVQERTIFF
(101) KDDGNYKTRAEVKFEGDTLVNRIEL
(126) KGIDFKEDGNILGHKLEYNYNSHNV
(151) YIMADKQKNGIKVNFKIRHNIEDGS
(176) VQLADHYQQNTPIGDGPVLLPDNHY
(201) LSYQSALSKDPNEKRDHMVLLEFVT
(226) AAGITLGMDELYK

**mCherry**

**MW = 30361 Da**
**$\varepsilon_{587nm}$ = 72000 $M^{-1}\cdot cm^{-1}$**

(Histag) MSYYHHHHHHDYDIPTTENLYFQGA
(Histag) MDPPVATM
(1) VSKGEEDNMAIIKEFMRFKVHMEGS
(26) VNGHEFEIEGEGEGRPYEGTQTAKL
(51) KVTKGGPLPFAWDILSPQFMYGSKA
(76) YVKHPADIPDYLKLSFPEGFKWERV
(101) MNFEDGGVVTVTQDSSLQDGEFIYK
(126) VKLRGTNFPSDGPVMQKKTMGWEAS
(151) SERMYPEDGALKGEIKQRLKLKDGG
(176) HYDAEVKTTYKAKKPVQLPGAYNVN
(201) IKLDITSHNEDYTIVEQYERAEGRH
(226) STGGMDELYK

*Scheme S1: Sequence of the 6 FPs. The region corresponding to the Histag is identical for all FPs except EGFP and mCherry, which contain 8 additional residues. For the five FPs derived from Aequora victoria (all except mCherry), residues that vary among these proteins are highlighted in bold. For each sequence, symbols shown below these positions indicate which proteins exhibit variations at the corresponding sites. The three residues involved in the chromophore structure are displayed in color.*

*Molecular weights (MW) were calculated using MassXpert software[1]. Molar absorption coefficients are from the FPbase database[2].*

### B) Quantitative assessment of kinetic parameters for FP photobleaching

#### B.1 Actometry measurements and instrumental specifications

##### Conversion of Photon Flux Density to Irradiance Values

Photon flux density $\boldsymbol{P}$ ($mol{\cdot}m^{-2}{\cdot}s^{-1}$) is the relevant parameter for the kinetic analysis of photobleaching. Irradiance, $\boldsymbol{I}$ ($W{\cdot}cm^{-2}$), commonly measured using a power meter, is nevertheless the parameter most often used by microscopists to compare experimental setups. The relationship between the two parameters is given by:

$$\boldsymbol{I} = \frac{\boldsymbol{h} \cdot \boldsymbol{c} \cdot \boldsymbol{N_A}}{\boldsymbol{10000} \cdot \boldsymbol{\lambda_{ex}}} \cdot \boldsymbol{P} \approx \frac{\boldsymbol{P}}{\boldsymbol{83333} \cdot \boldsymbol{\lambda_{ex}}}$$

with $\boldsymbol{h}$ the Planck constant ($6.63{\cdot}10^{-34}$ $m^2{\cdot}kg{\cdot}s^{-1}$), $\boldsymbol{c}$ the speed of light in vacuum ($3.00{\cdot}10^{8}$ $m{\cdot}s^{-1}$), $\boldsymbol{N_A}$ the Avogadro number ($6.02{\cdot}10^{23}$ $mol^{-1}$) and $\boldsymbol{\lambda_{ex}}$ the wavelength of the source (m).

The measurements of both $\boldsymbol{P}$ and $\boldsymbol{I}$ undergone by samples during photobleaching experiments were determined using Dronpa2 photoswitching as an actinometer, as described in Lahlou et al. [3,4].

##### Wide field epifluorescence pathway of the microscope

Purified His-tagged Dronpa2 FPs were first immobilized on Ni-NTA agarose beads using the same protocol as for photobleaching experiments (see Methods). The Dronpa2-decorated beads were then illuminated using either the GFP or YFP filter sets, at 0.95% and 1.2% of the maximum irradiance, respectively, in order to obtain phototoswitching kinetics slow enough to be accurately recorded (Figure S1a). To ensure irradiance linearity at such low output levels, neutral density filters with transmittance of 9.5% and 12% respectively were used with an input lamp power of 10%, a value sufficiently high to ensure measurements in the linearity range of the lamp intensity. The emission spectra of the lamp for each spectral selection were measured using a fiber-coupled spectrophotometer (Ocean Optics SR6). These spectral profiles were combined with (i) the monoexponential time constants computed from kinetics in figure S1a, (ii) the excitation spectrum of Dronpa2 (extracted from online database associated with ref. [3]) and (iii) its tabulated photoswitching molar cross-sections at 480 nm and 500 nm[3]. This approach allowed reconstruction of the scaled emission spectra of the light source at the sample position for an input lamp power at 100 % (figure S1b) according to the computing method described by Lahlou et al[3]. Finally, the absorption spectra of studied FPs (measured on the BEAM setup) and the scaled emission spectra of the light source were used to compute the equivalent monochromatic photon flux at the absorption maximum of each FP (table S1). The detailed model is presented below.

The kinetics of the Dronpa2 photoswitching reaction, monitored through the resulting decrease in fluorescence intensity, follows a monoexponential decay (see Figures S1a and S2a). Under monochromatic excitation, the rate constant obtained with this model, $\boldsymbol{k_{switch}}$, is linked to photon flux density $\boldsymbol{P}$ by:

$$\boldsymbol{k_{switch}}(\boldsymbol{\lambda_{ex}}) = \boldsymbol{\sigma_{switch}}(\boldsymbol{\lambda_{ex}}) \cdot \boldsymbol{P}(\boldsymbol{\lambda_{ex}})$$

Where $\boldsymbol{\lambda_{ex}}$ **is** the excitation wavelength and $\boldsymbol{\sigma_{switch}}$ **is** the photoswitching molar cross section of Dronpa2.

Under polychromatic excitation, the rate constant becomes:

$$\boldsymbol{k_{switch}} = \int_{\lambda_{min}}^{\lambda_{max}} \boldsymbol{\sigma_{switch}(\lambda) \cdot p(\lambda) \cdot d\lambda}$$

where is $\boldsymbol{p(\lambda)}$ the spectral photon flux density (mol·m$^{-2}$·s$^{-1}$·nm$^{-1}$), and $\boldsymbol{\lambda_{min}}$ and $\boldsymbol{\lambda_{max}}$ the wavelengths delimiting the range of the light source and the excitation spectrum of Dronpa2.
$\boldsymbol{\sigma_{switch}(\lambda)}$ was estimated from the two tabulated values reported in ref. [3], $\boldsymbol{\sigma_{switch}(480)}$ = 198 mol·m$^{-2}$·s$^{-1}$ and $\boldsymbol{\sigma_{switch}(500)}$ = 128 mol·m$^{-2}$·s$^{-1}$, assuming that it is locally proportional (within ± 15 nm) to the absorption coefficient of Dronpa2 $\boldsymbol{\varepsilon_D(\lambda)}$ whose relative value was obtained from the absorption spectrum available in the database with ref. [3]. Accordingly:

$$\boldsymbol{\sigma_{switch}(\lambda) = \sigma_{switch}(\lambda_{ref}) \frac{\varepsilon_D(\lambda)}{\varepsilon_D(\lambda_{ref})}}$$

With $\boldsymbol{\lambda_{ref}}$ = 480 or 500 nm.
The purpose is then to determine $\boldsymbol{p(\lambda)}$ over the full spectral range, which corresponds to the scaled spectrum of the light source. To this end, a scaling factor $\boldsymbol{S}$ is introduced to relate $\boldsymbol{p(\lambda)}$ to $\mathbf{L_{norm}}(\boldsymbol{\lambda})$, the emission spectrum of the polychromatic lamp normalized to unit area.

$$\boldsymbol{p(\lambda) = S \cdot L_{norm}(\lambda)}$$

$$\boldsymbol{L_{norm}(\lambda) = \frac{L(\lambda)}{\int_{\lambda_{min}}^{\lambda_{max}} L(\lambda) \cdot d\lambda}}$$

with $\mathbf{L}(\boldsymbol{\lambda})$ the measured spectrum of the polychromatic source. Next, we introduce the action spectrum of photoswitching, $\boldsymbol{AS(\lambda)}$, which accounts for the overlap between the photoswitching spectrum of Dronpa2 and the emission spectrum of the polychromatic source. This spectrum is obtained as follows:

$$\boldsymbol{AS(\lambda) = \sigma_{switch}(\lambda) \cdot L_{norm}(\lambda)}$$

We can now link the integral of the action spectrum $\{\boldsymbol{AS}\}$ to $\boldsymbol{k_{switch}}$:

$$\boldsymbol{k_{switch}} = \int_{\lambda_{min}}^{\lambda_{max}} \boldsymbol{\sigma_{switch}(\lambda) \cdot S \cdot L_{norm}(\lambda) \cdot d\lambda = S \cdot} \int_{\lambda_{min}}^{\lambda_{max}} \boldsymbol{AS(\lambda) \cdot d\lambda = S \cdot \{AS\}}$$

where $\boldsymbol{S}$ is a scaling factor. This factor can be determined from the experimentally measured $\boldsymbol{k_{switch}}$, and the integrated action spectrum, $\{\boldsymbol{AS_{norm}}\}$, obtained from spectral measurements. Finally, $\boldsymbol{p(\lambda)}$ is computed as follows:

$$\boldsymbol{p(\lambda) = \frac{k_{switch}}{\{AS\}} \cdot L_{norm}(\lambda)}$$

The scaling factors $\boldsymbol{S_{GFP}}$ and $\boldsymbol{S_{YFP}}$ were experimentally determined on our setup for excitation with the GFP and YFP filter cubes, respectively (see the corresponding kinetics in Figure S1a and setup specifications in Table S1), leading to similar values (relative difference < 5%). For CFP and RFP filter cubes, for which no Dronpa2 photoswitching kinetics can be performed, we used the average scaling value $\langle \boldsymbol{S} \rangle = \frac{\boldsymbol{S_{GFP}}+\boldsymbol{S_{YFP}}}{2}$. Finally, the scaled spectra $\boldsymbol{p(\lambda)}$ were obtained for excitation with all four configurations of our setup (Figure S1b).

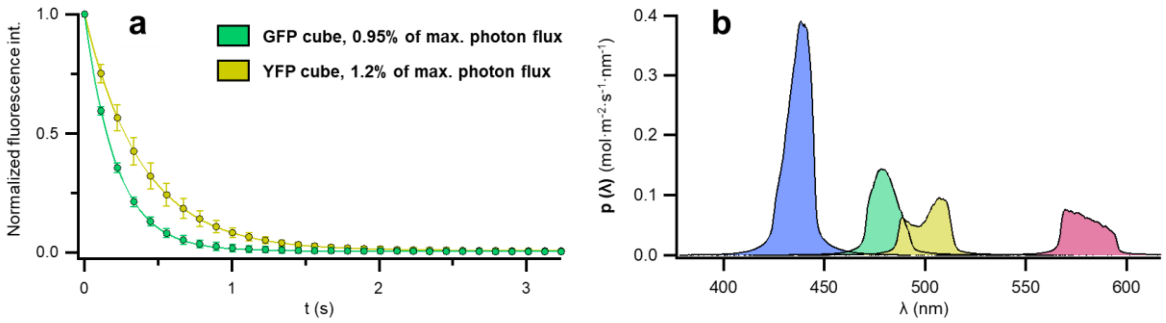

*Figure S1: a) Photoswitching kinetics of Dronpa2 immobilized on Ni-TNA agarose beads and illuminated via the epifluorescence pathway of the microscope. Each curve represents the average of 10 photobleaching experiments with error bars indicating the standard deviation. Monoexponential fits are shown as solid lines. b) Scaled emission spectra of the epifluorescence lamp used for photobleaching experiments with CFP, GFP, YFP, or RFP filter cubes (blue, green, yellow, and red, respectively). The scaling corresponds to the maximum input power of the LED source*

Next, for each studied FP, we calculated the equivalent monochromatic photon flux density $\boldsymbol{P^{cal}_{\lambda_{max}}}$, corresponding to the photon flux delivered by the light source at the wavelength of maximum of FP absorption $\boldsymbol{\lambda_{max}}$ (see table S1).

$$\boldsymbol{P^{cal}_{\lambda_{max}}} = \int_{\lambda_{min}}^{\lambda_{max}} \boldsymbol{p(\lambda)} \cdot \frac{\boldsymbol{\varepsilon_{FP}(\lambda)}}{\boldsymbol{\varepsilon_{FP}(\lambda_{max})}} \cdot \boldsymbol{d\lambda}$$

Where $\boldsymbol{\varepsilon_{FP}}$ is the molar absorption coefficient of the considered FP. The value of $\boldsymbol{P^{cal}_{\lambda_{max}}}$ was then used to determine the photobleaching molar cross section at $\boldsymbol{\lambda_{max}}$, $\boldsymbol{\sigma^{bleach}_{\lambda_{max}}}$, for each FP using an equation similar to the monochromatic equation introduced earlier:

$$\boldsymbol{\sigma^{bleach}_{\lambda_{max}}} = \frac{\boldsymbol{k_{bleach}}}{\boldsymbol{P^{cal}_{\lambda_{max}}}}$$

With $\boldsymbol{k_{bleach}}$ the photobleaching rate constant obtained from photobleaching kinetics measurements. The results of photobleaching experiment performed using the microscope setup are presented in section B.2.

| FP | cube | Excitation filter | Dichroic mirror | Emission filter | Total photon Flux density (mol·m$^{-2}$·s$^{-1}$) | $P_{\lambda_{max}}^{cal}$ equivalent photon flux density at $\lambda_{max}$ (mol·m$^{-2}$·s$^{-1}$) |
|---|---|---|---|---|---|---|
| Aquamarine | CFP (CFP 2432C) | FF02-438/24 | FF458-Di02 | FF01-483/32 | 6.24 | 5.89 [162.88]* |
| mTurquoise | CFP (CFP 2432C) | FF02-438/24 | FF458-Di02 | FF01-483/32 | 6.24 | 5.93 [164.04]* |
| EGFP | GFP (GFP 1828A) | FF02-482/18 | FF495-Di03 | FF02-520-28 | 2.55 | 2.26 [55.63]* |
| Citrine | YFP (YFP 2427B) | FF01-500/24 | FF520-Di02 | FF01-542/27 | 2.02 | 1.25 [29.10]* |
| EYFP | YFP (YFP 2427B) | FF01-500/24 | FF520-Di02 | FF01-542/27 | 2.02 | 1.30 [30.22]* |
| mCherry | RFP (C156423 custom) | ET-580/25 | ZT594rdc | ET-625/30 | 1.77 | 1.53 [31.20]* |

*Table S1: Specifications of the filter sets and corresponding photon flux density (mol·m$^{-2}$·s$^{-1}$) for epifluorescence pathway of the microscope. Values are reported at the maximum input power of the LED source. * corresponding irradiance (W·cm$^{-2}$)*

#### BEAM setup

The effective photon flux delivered to a 35 µL solution volume contained in a 0.3x0.3 cm cuvette within the BEAM custom-built setup was determined as previously described[4]. Photon flux (mol·m$^{-2}$·s$^{-1}$) or irradiance (W·cm$^{-2}$) of the 445 nm laser diode was measured using Dronpa2 photoswitching kinetics (figure S2a), whereas that of the 515 nm laser diode was quantified via ratiometric fluorescence intensity measurement of the FP mNeongreen (Figure S2b).

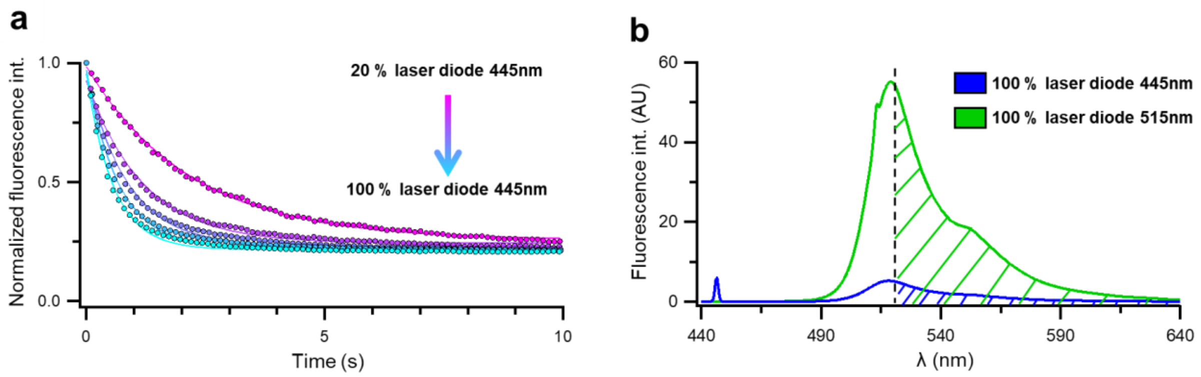

*Figure S2: Actinometry measurements for the two laser diodes of the BEAM setup. a) Photoswitching kinetics of Dronpa2 in a 35 µL solution contained in a 0.3x0.3 cm cuvette and illuminated with the 445 nm laser diode used for in vitro photobleaching experiments. The monoexponential fits are the solid lines. b) Comparison of the emission spectra of mNeonGreen excited with the 445 nm or the 515 nm laser diode. The intensity ratio was calculated by integrating the signal over the hatched regions.*

All photobleaching experiments were conducted at the maximum input power of each laser source, corresponding to irradiances of 0.39 W·cm$^{-2}$ at 445 nm and a 0.60 W·cm$^{-2}$ at 515 nm. To account for the fact that the excitation wavelength of the laser diode ($\lambda_{ex}$) does not necessarily coincide with the absorption maximum ($\lambda_{max}$) of the FP, the equivalent photon flux density at $\lambda_{max}$ was calculated for each FP according to :

$$P_{\lambda_{max}}^{cal} = P_{\lambda_{ex}}^{mes} \cdot \frac{\varepsilon_{FP}(\lambda_{ex})}{\varepsilon_{FP}(\lambda_{max})}$$

With $P_{\lambda_{ex}}^{mes}$ the measured photon flux density at the excitation wavelength and $P_{\lambda_{max}}^{cal}$ the equivalent photon flux density at $\lambda_{max}$.

| FP | $\lambda_{ex}$ laser (nm) | $P_{\lambda_{ex}}^{mes}$ (mol·m$^{-2}$·s$^{-1}$) | $\lambda_{max}$ FP (nm) | $\frac{\varepsilon_{\lambda_{ex}}}{\varepsilon_{\lambda_{max}}}$ | $P_{\lambda_{max}}^{cal}$ (mol·m$^{-2}$·s$^{-1}$) |
|---|---|---|---|---|---|
| Aquamarine | 445 | $1.44 \cdot 10^{-2}$ [0.39]* | 430 | 0.92 | $1.32 \cdot 10^{-2}$ [0.37]* |
| mTurquoise | 445 | $1.44 \cdot 10^{-2}$ [0.39]* | 434 | 0.93 | $1.34 \cdot 10^{-2}$ [0.37]* |
| EGFP | 445 | $1.44 \cdot 10^{-2}$ [0.39]* | 488 | 0.39 | $5.61 \cdot 10^{-3}$ [0.14]* |
| Citrine | 515 | $2.57 \cdot 10^{-2}$ [0.60]* | 515 | 1 | $2.57 \cdot 10^{-2}$ [0.60]* |
| EYFP | 515 | $2.57 \cdot 10^{-2}$ [0.60]* | 515 | 1 | $2.57 \cdot 10^{-2}$ [0.60]* |
| mCherry | 515 | $2.57 \cdot 10^{-2}$ [0.60]* | 580 | 0.20 | $5.13 \cdot 10^{-3}$ [0.11]* |

*Table S2: Photon flux density (mol·m$^{-2}$·s$^{-1}$) specifications for BEAM photobleaching experiments. * corresponding irradiance (W·cm$^{-2}$)*

#### B.2 Photobleaching kinetics of FPs

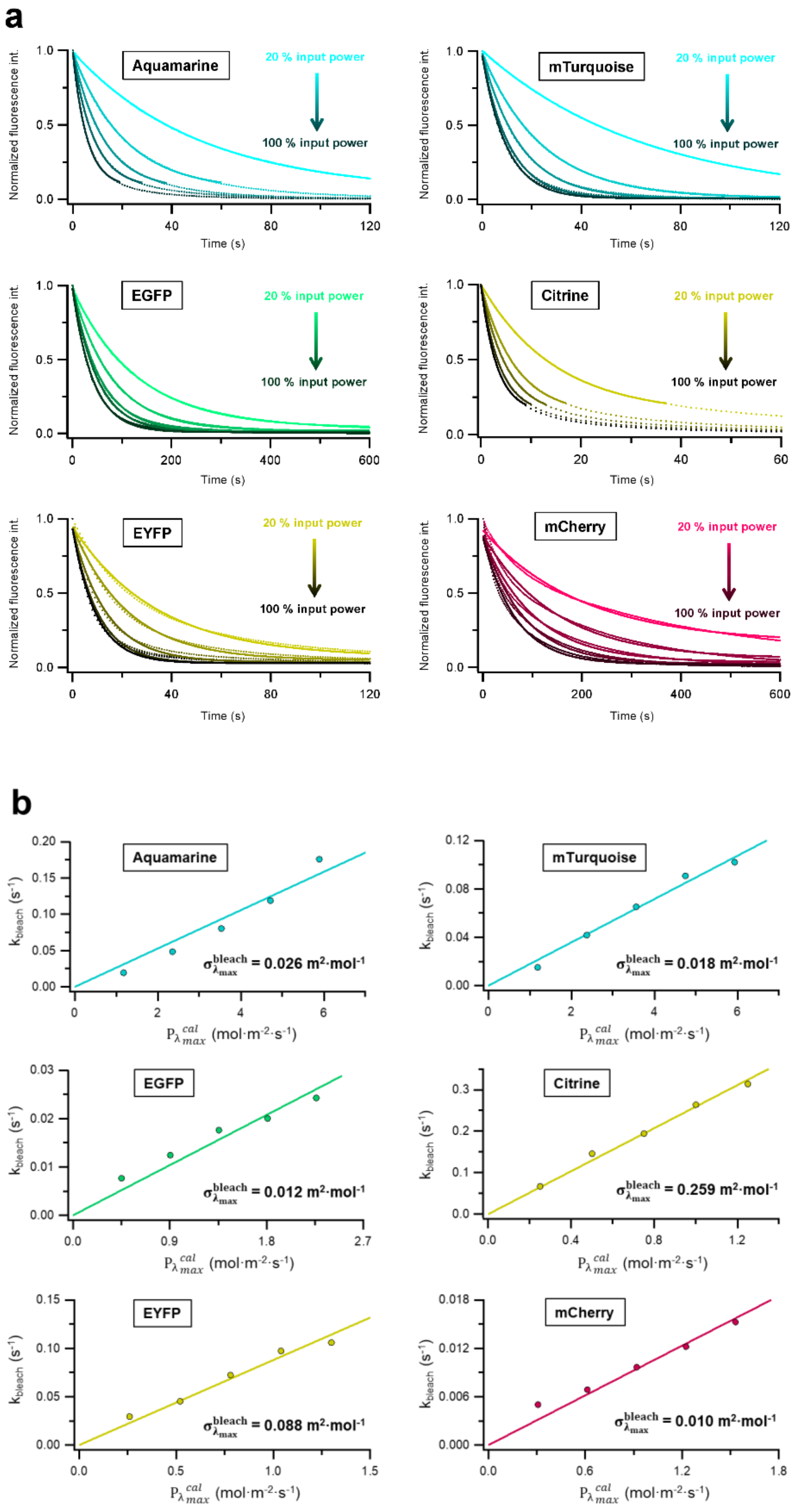

*Figure S3: (a) Photobleaching kinetics of FPs* ***in living COS-7 cells*** *acquired using the epifluorescence pathway of the microscope at various input powers. Experimental data are shown as dots, with corresponding monoexponential fits displayed as solid lines. The decays of fluorescence intensity of Aquamarine and Citrine exhibit biexponential behavior at long bleaching times. To restrict the analysis to the monoexponential regime, fits were truncated at 10 % and 20 % of the remaining fluorescence intensity, respectively. (b) Evolution of the photobleaching kinetic rate constants determined in (a) with the equivalent photon flux density calculated at each FP absorption maximum. Linear fits are shown in solid lines. The slopes, corresponding to the photobleaching molar cross-sections of the FPs, are indicated.*

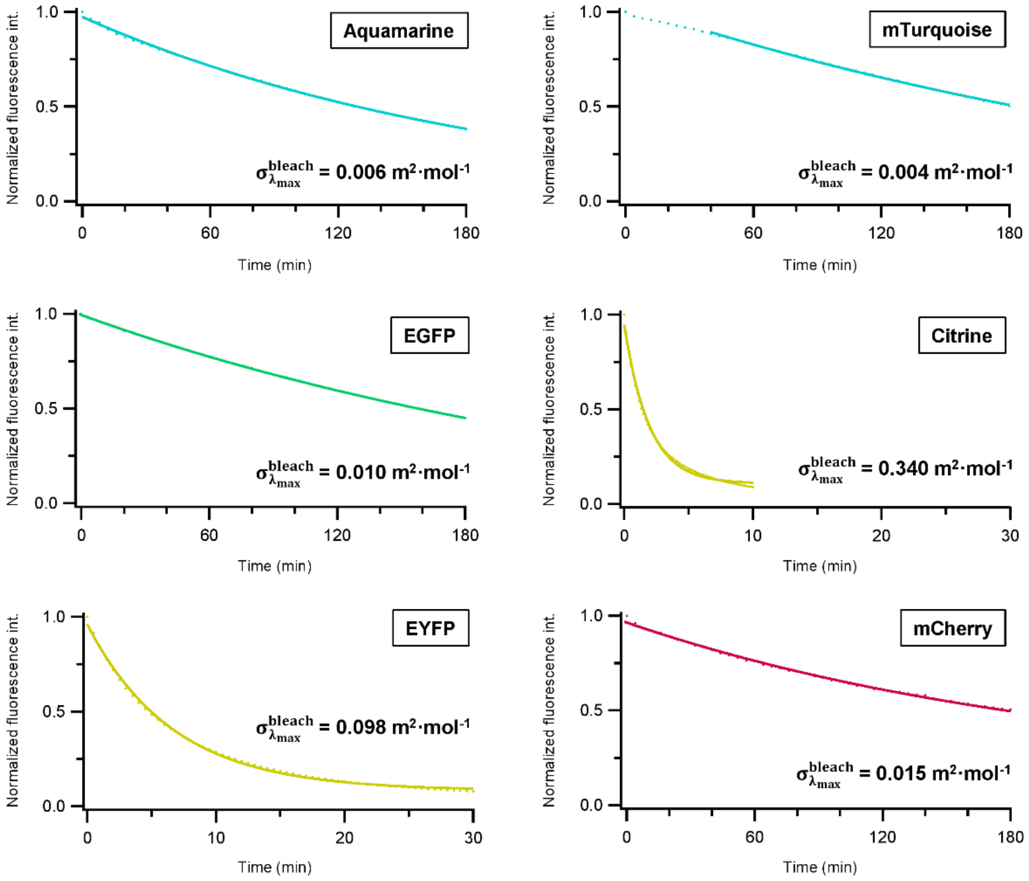

*Figure S4: Photobleaching kinetics of* ***purified FPs in solution****, acquired using the BEAM setup. Sample was continuously illuminated in a 0.3x0.3 cm cuvette (0.35 µL of FP) using either a 445 nm or a 515 nm laser diode. FP concentrations were adjusted to yield an absorbance of approximately 0.1 at the excitation wavelength, thereby optimizing the fluorescence signal while minimizing the inner filter effect. Monoexponential fits are displayed in solid lines. For mTurquoise, a transient regime was observed at the beginning of its photobleaching kinetics, therefore, only data points collected after 40 min of illumination were used for the fit.*

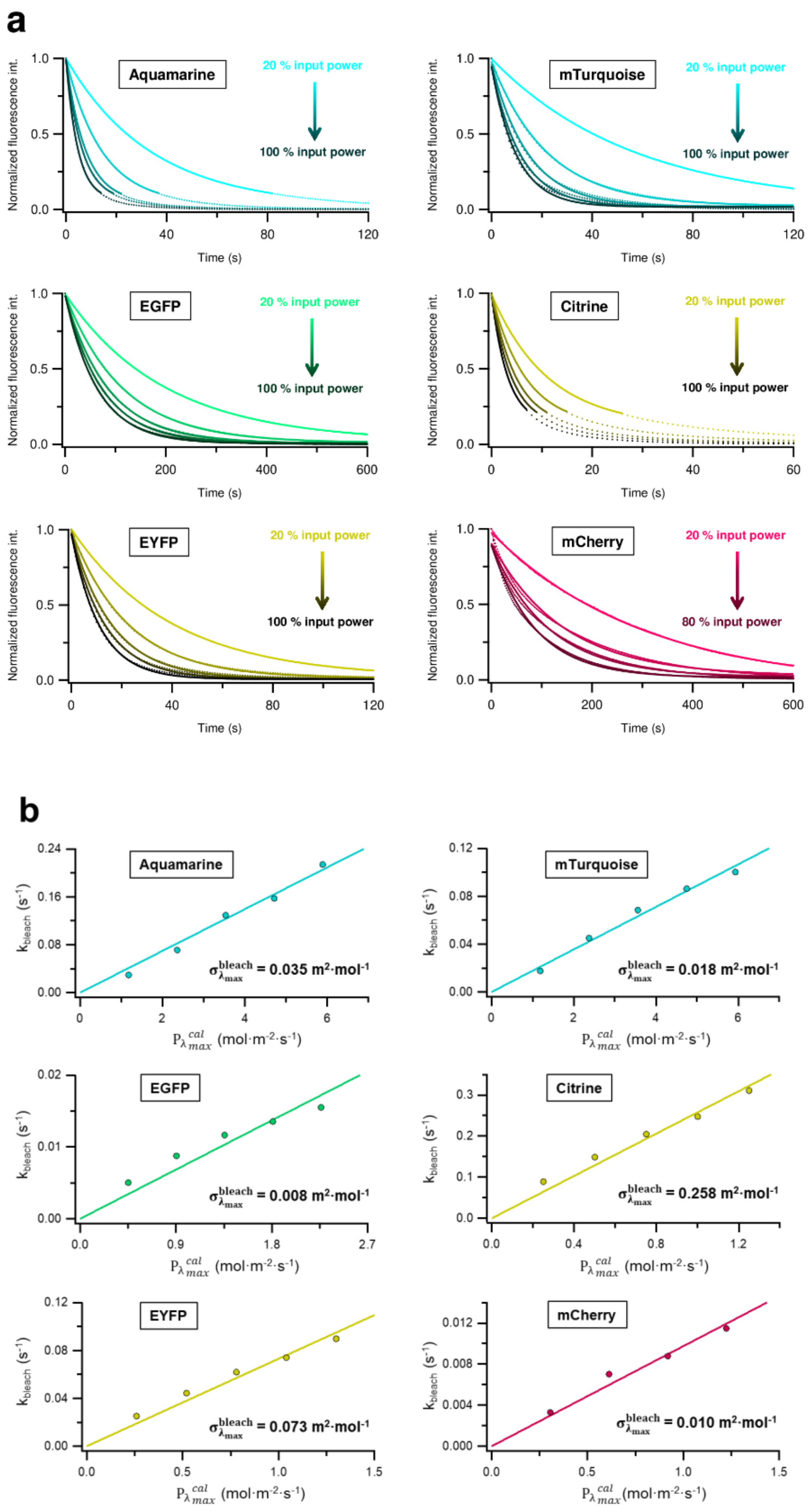

*Figure S5: (a). Photobleaching kinetics of* ***purified FPs bound to Ni-NTA beads****, acquired using the epifluorescence pathway of the microscope at various input powers. Experimental curves are shown as dots, with corresponding monoexponential fits displayed as solid lines. The decays of fluorescence intensity of Aquamarine and Citrine exhibit biexponential behavior at long bleaching times. To restrict the analysis to the monoexponential regime, fits were truncated at 10 % and 20 % of the remaining fluorescence intensity, respectively. (b) Evolution of the photobleaching kinetic rate constants determined in (a) with the equivalent photon flux density calculated at each FP's absorption maximum. Linear fits are shown in solid lines. The slopes, corresponding to the photobleaching molar cross-sections of the FPs, are indicated.*

| | $\sigma_{\lambda_{max}}^{bleach}$ ($m^2 \cdot mol^{-1}$) | | | Relative $\sigma_{\lambda_{max}}^{bleach}$ (%) | | | $\Phi_{bleach}$ | | | Relative $\Phi_{bleach}$ (%) | | |
|---|---|---|---|---|---|---|---|---|---|---|---|---|
| **Protein** | In vitro | Agarose beads | In cellulo | In vitro | Agarose beads | In cellulo | In vitro | Agarose beads | In cellulo | In vitro | Agarose beads | In cellulo |
| **Aquamarine** | $6.4 \cdot 10^{-3}$ | $3.5 \cdot 10^{-2}$ | $2.6 \cdot 10^{-2}$ | 1.9 | 13.5 | 10.2 | $1.1 \cdot 10^{-6}$ | $5.8 \cdot 10^{-6}$ | $4.4 \cdot 10^{-6}$ | 5.8 | 41.6 | 31.4 |
| **mTurquoise** | $3.9 \cdot 10^{-3}$ | $1.8 \cdot 10^{-2}$ | $1.8 \cdot 10^{-2}$ | 1.2 | 6.9 | 6.9 | $5.6 \cdot 10^{-7}$ | $2.6 \cdot 10^{-6}$ | $2.6 \cdot 10^{-6}$ | 3.1 | 18.4 | 18.4 |
| **EGFP** | $1.0 \cdot 10^{-2}$ | $7.6 \cdot 10^{-3}$ | $1.2 \cdot 10^{-2}$ | 3.1 | 3.0 | 4.4 | $8.0 \cdot 10^{-7}$ | $5.9 \cdot 10^{-7}$ | $9.0 \cdot 10^{-7}$ | 4.4 | 4.2 | 6.4 |
| **Citrine** | $3.4 \cdot 10^{-1}$ | $2.6 \cdot 10^{-1}$ | $2.6 \cdot 10^{-1}$ | 100 | 100 | 100 | $1.8 \cdot 10^{-5}$ | $1.4 \cdot 10^{-5}$ | $1.4 \cdot 10^{-5}$ | 100 | 100 | 100 |
| **EYFP** | $9.8 \cdot 10^{-2}$ | $7.3 \cdot 10^{-2}$ | $8.8 \cdot 10^{-2}$ | 29.0 | 28.3 | 33.9 | $6.1 \cdot 10^{-6}$ | $4.5 \cdot 10^{-6}$ | $5.4 \cdot 10^{-6}$ | 33.2 | 32.4 | 38.7 |
| **mCherry** | $1.5 \cdot 10^{-2}$ | $9.8 \cdot 10^{-3}$ | $1.0 \cdot 10^{-2}$ | 4.6 | 3.8 | 4.0 | $9.3 \cdot 10^{-7}$ | $5.9 \cdot 10^{-7}$ | $6.2 \cdot 10^{-7}$ | 5.1 | 4.2 | 4.4 |

*Table S3: Photobleaching characteristics of FPs under different conditions (in vitro, agarose beads and in live cells). Photobleaching cross sections ($\boldsymbol{\sigma_{\lambda_{max}}^{bleach}}$), corresponding quantum yields ($\boldsymbol{\phi_{bleach}}$), and their relative values are reported. Data correspond to those shown in Figure 1.*

#### C) Fluorescence lifetimes decrease with photobleaching

##### FLIM measurements of FPs expressed in COS-7 cells

| FP | Excitation wavelength | Filter1 | Filter2 | Filter3 |
|---|---|---|---|---|
| Aquamarine | 440 nm | > 458 nm | > 458 nm | 480 ± 15 nm |
| mTurquoise | 440 nm | > 458 nm | > 458 nm | 480 ± 15 nm |
| EGFP | 466 nm | > 488 nm | | 535 ± 20 nm |
| Citrine | 466 nm | > 488 nm | | 535 ± 20 nm |
| EYFP | 466 nm | > 488 nm | | 535 ± 20 nm |

*Table S4: Excitation light sources and emission filters used in the TCSPC pathway of the microscope.*

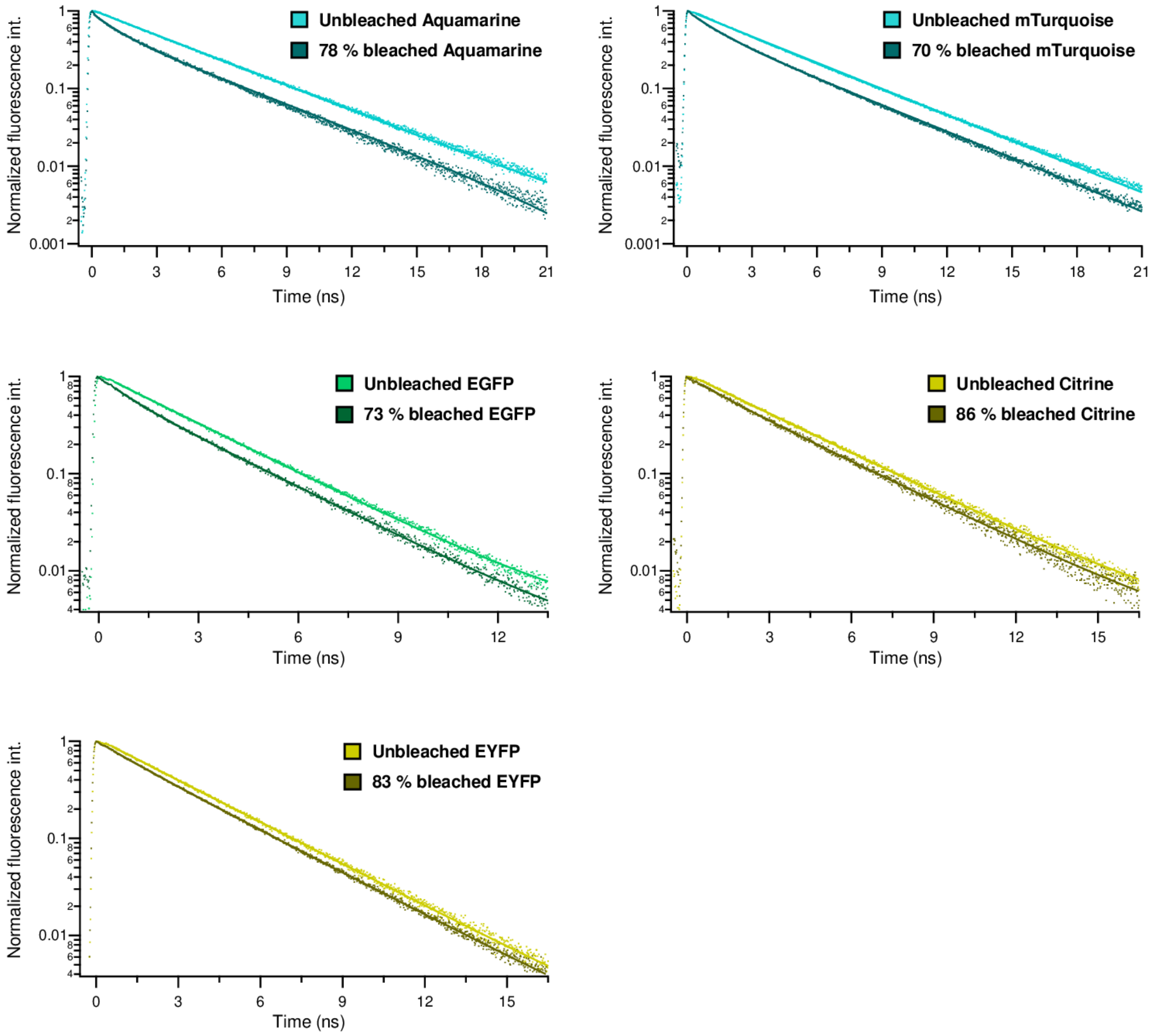

*Figure S6: Examples of fluorescence decay curves of **FPs expressed in COS-7 cells**, acquired by FLIM. Each panel displays decay curve from multiple cells within the field of view, recorded both before and after photobleaching. Experimental data are shown as dots and fitted curves as solid lines. A monoexponential model was used for unbleached FPs, whereas a biexponential model was applied to bleached FPs, with the long lifetime component fixed to the lifetime value measured for the corresponding unbleached FP (see Methods).*

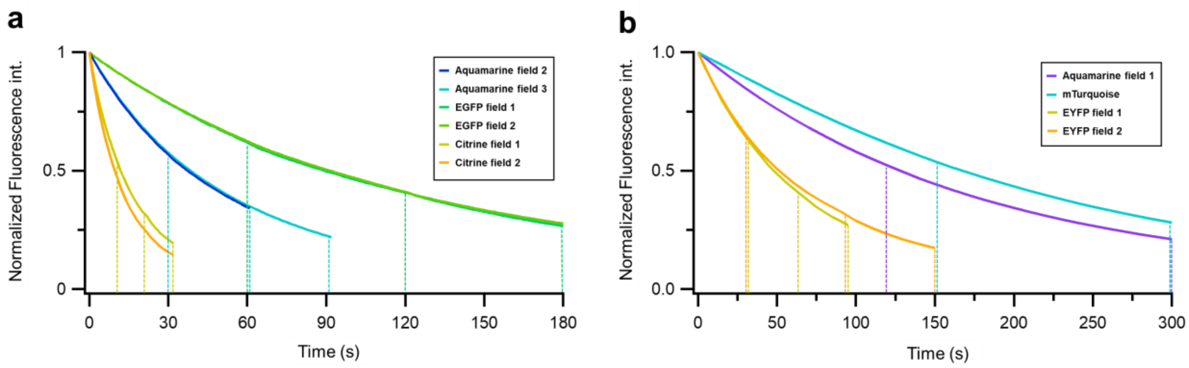

*Figure S7: Photobleaching kinetics of fluorescent proteins expressed in COS-7 cells. Vertical dashed lines indicate time points at which bleaching was paused to acquire FLIM images. a) Experiments performed at 20°C using 10% lamp power. b) Experiments performed at 26°C with 20% lamp power. Corresponding fluorescence lifetime values are presented in Figure 2 and representative decay curves are displayed in Figure S6.*

#### Fluorescence Lifetime measurements of FPs in vitro

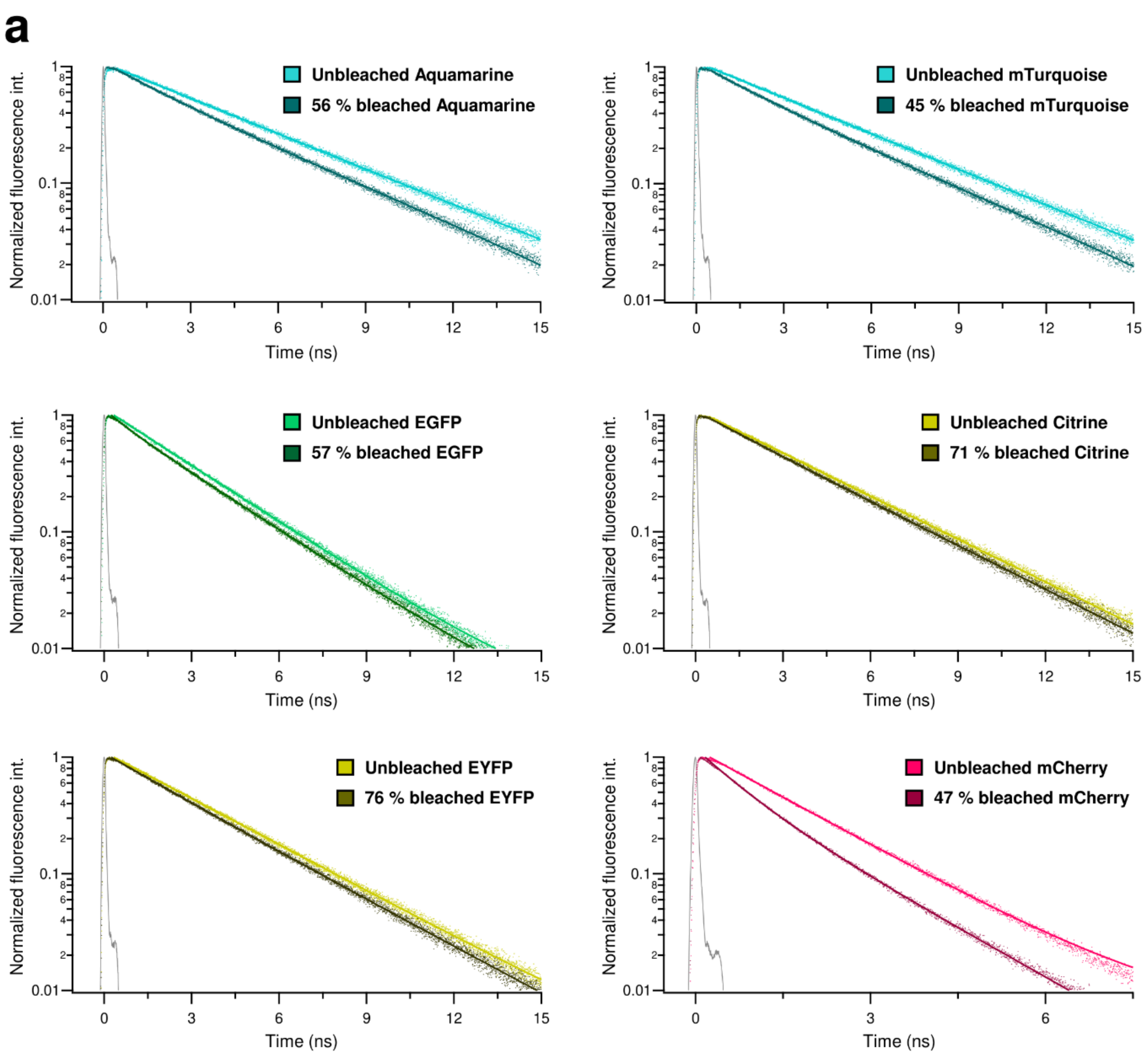

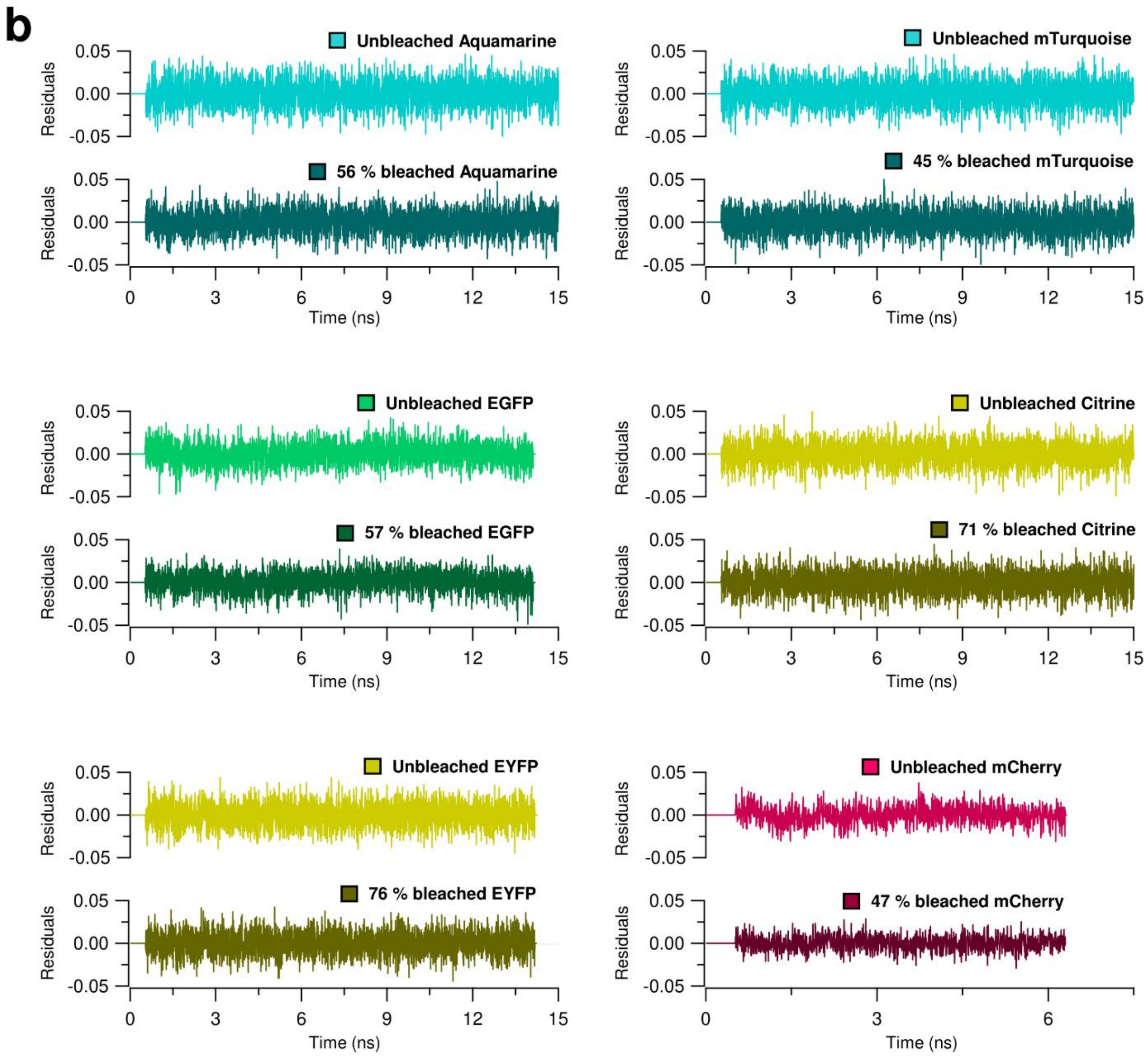

*Figure S8: a) Representative fluorescence decay curves of **purified FPs in solution**, acquired before and after photobleaching using laser diodes. FP solutions were photobleached using the BEAM setup and subsequently diluted to maintain an absorbance below 0.1, ensuring accurate measurements of the fluorescence decay curve. Experimental data are shown as dots and fitted curves as solid lines. A mono-exponential model was used for unbleached FPs, whereas a biexponential model was applied to bleached FPs, with the long lifetime component fixed to the lifetime value measured for the corresponding unbleached FP (see Methods). b) Residuals of the corresponding fits.*

| FP | $\tau_0$ (ns) | $\tau_{damaged}$ (ns) | Final $\langle\tau\rangle$ (ns) | $100 \cdot \frac{\tau_{damaged}}{\tau_0}$ (%) | $100 \cdot \alpha_{damaged}$ (%) |
|---|---|---|---|---|---|
| **Aquamarine** | 4.29 | 2.07 | 3.66 | 48 | 28 |
| **mTurquoise** | 4.22 | 2.03 | 3.63 | 48 | 27 |
| **EGFP** | 2.67 | 0.8 | 2.51 | 30 | 9 |
| **Citrine** | 3.53 | 1.97 | 3.37 | 56 | 10 |
| **EYFP** | 3.28 | 2.1 | 3.1 | 64 | 15 |
| **mCherry** | 1.58 | 0.7 | 1.23 | 44 | 40 |

*Table S5: Fluorescence lifetimes and fractions of damaged species formed upon photobleaching of each FP in solution. Lifetimes were estimated from the fits shown in Figure S8. Definitions of all parameters are provided in Sections E.4 and E.5.*

#### D) Spectroscopic study of FPs photobleaching

##### Calibration for inner filter effect correction

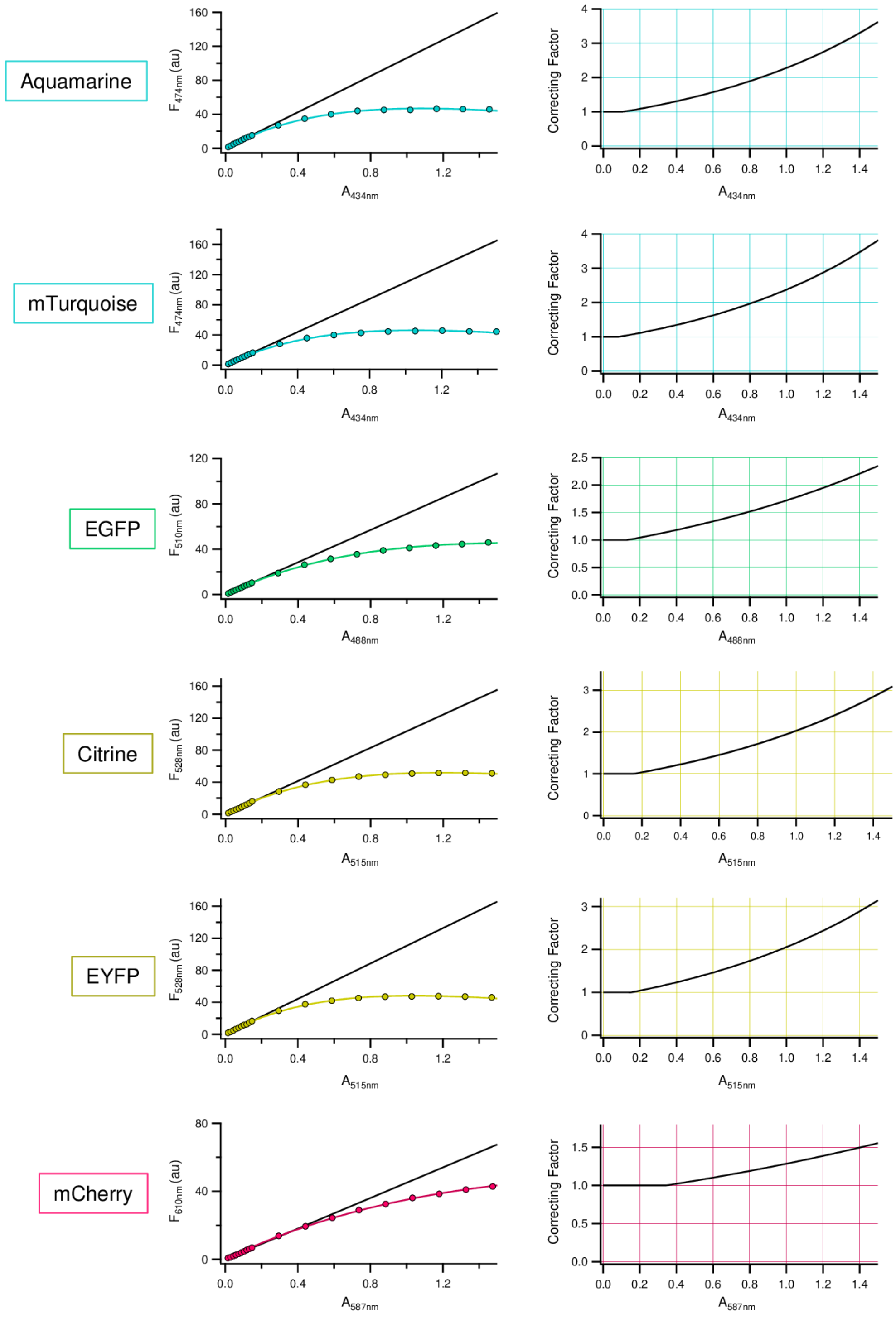

*Figure S9: Calibration curves for correction of the inner filter effect. FPs were excited under the conditions described in Tables S1 and S2 using the BEAM setup. The correction method is detailed in ref [4].*

#### Normalized absorption and fluorescence emission spectra

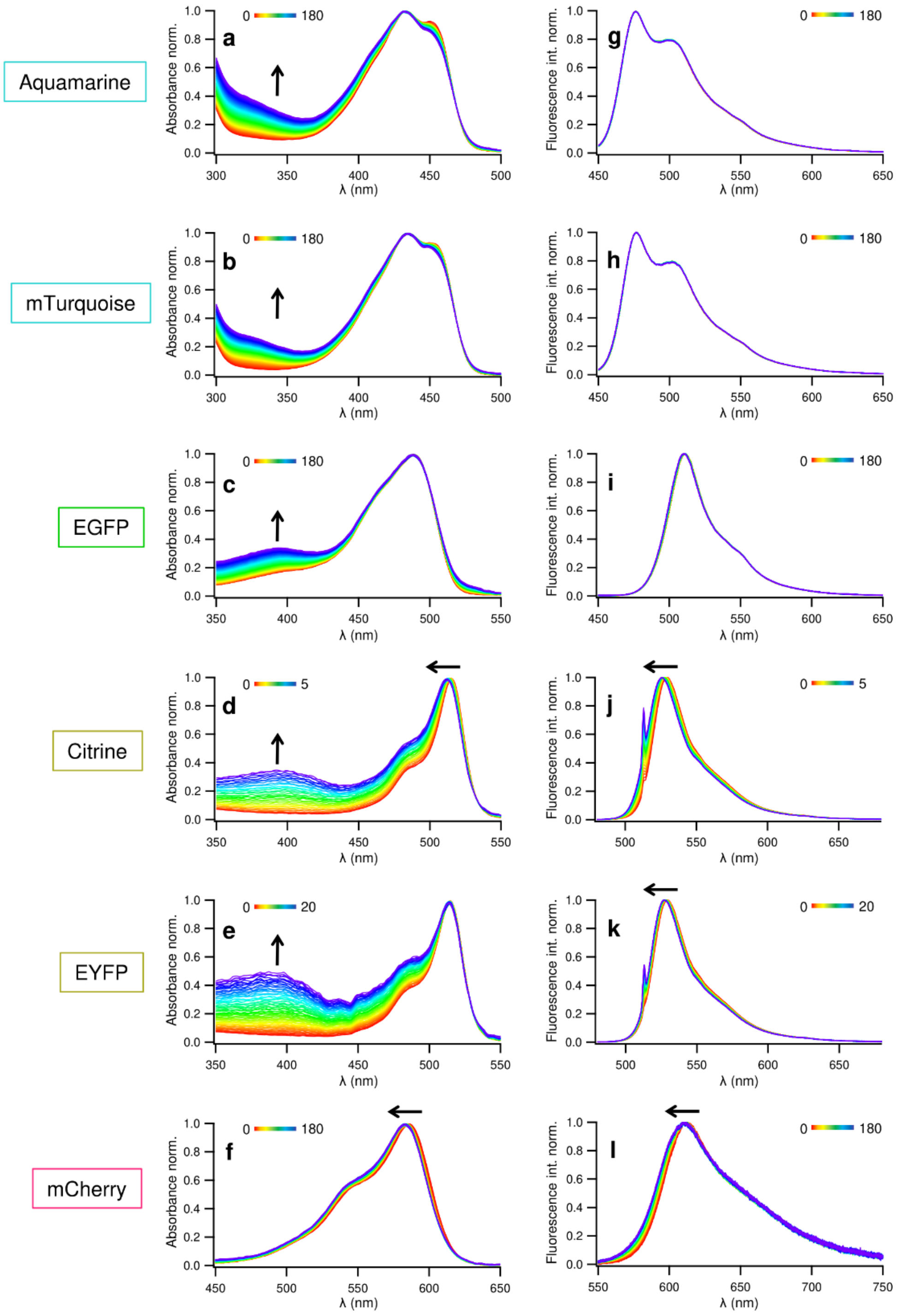

*Figure S10: Time-dependent changes of normalized absorption spectra (a-f) and normalized fluorescence emission spectra (g-l) under irradiation. The corresponding unnormalized spectra are shown in Figure 3. The initial maximum absorption of the FP solutions was set to* ***0.5****. Each row presents results obtained for the indicated FP (left). Irradiations were carried out using laser diodes at an irradiance of 0.4 W·cm$^{-2}$ at 445 nm for CFPs and EGFP, and 0.6 W·cm$^{-2}$ at 515 nm for YFPs and mCherry.*

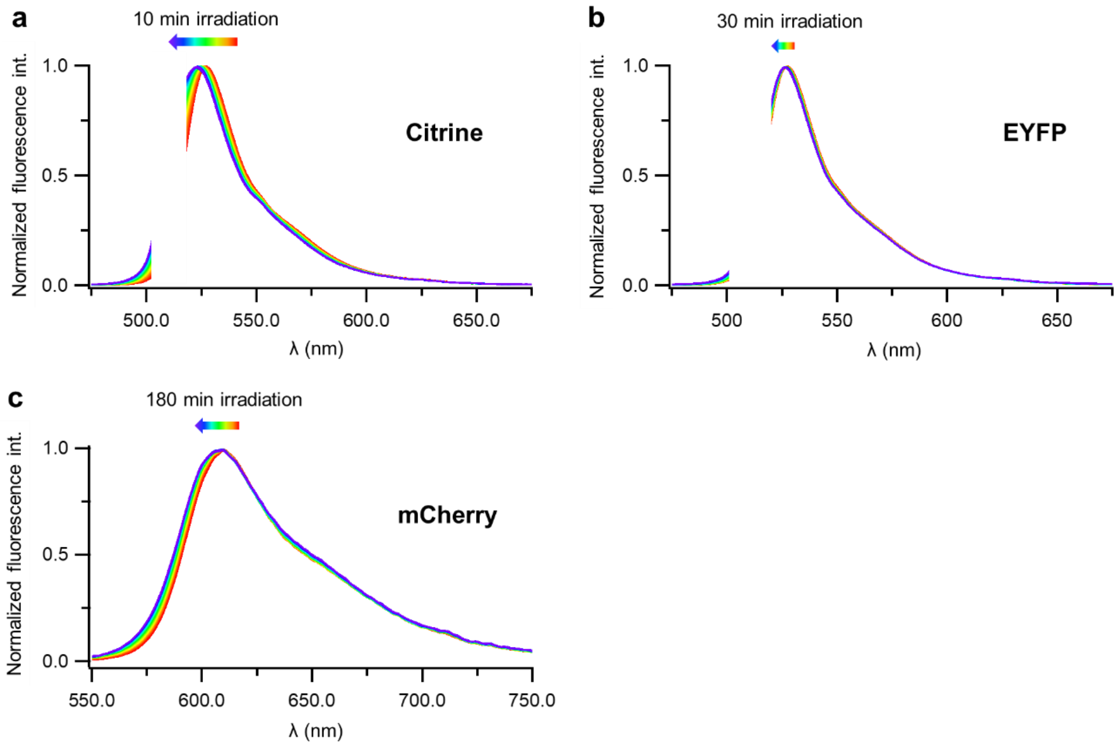

*Figure S11: Time-dependent changes of fluorescence emission spectrum under irradiation for (a) Citrine, (b) EYFP and (c) mCherry. The initial maximum absorption of the FP solutions was set to **0.1**. For Citrine and EYFP, the Rayleigh peaks of the 515 nm laser are particularly intense at this concentration and have been hidden for clarity. At this absorbance, no inner filter effect is present. The observed blue-shift for Citrine and mCherry arises from photobleaching, whereas EYFP exhibits a very slight shift.*

#### Photobleaching with white light source

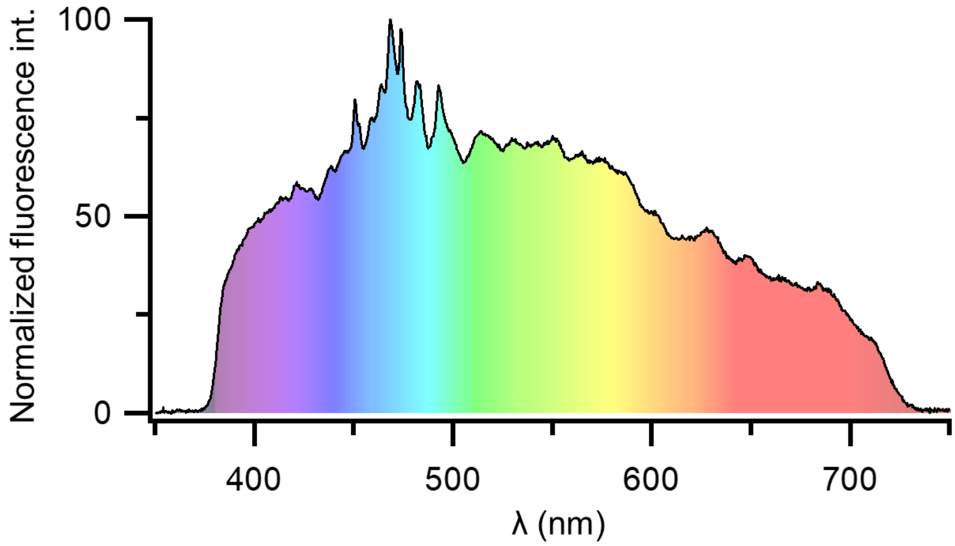

*Figure S12: Emission spectrum of the white light source (Eurosep) used for in vitro photobleaching experiments.*

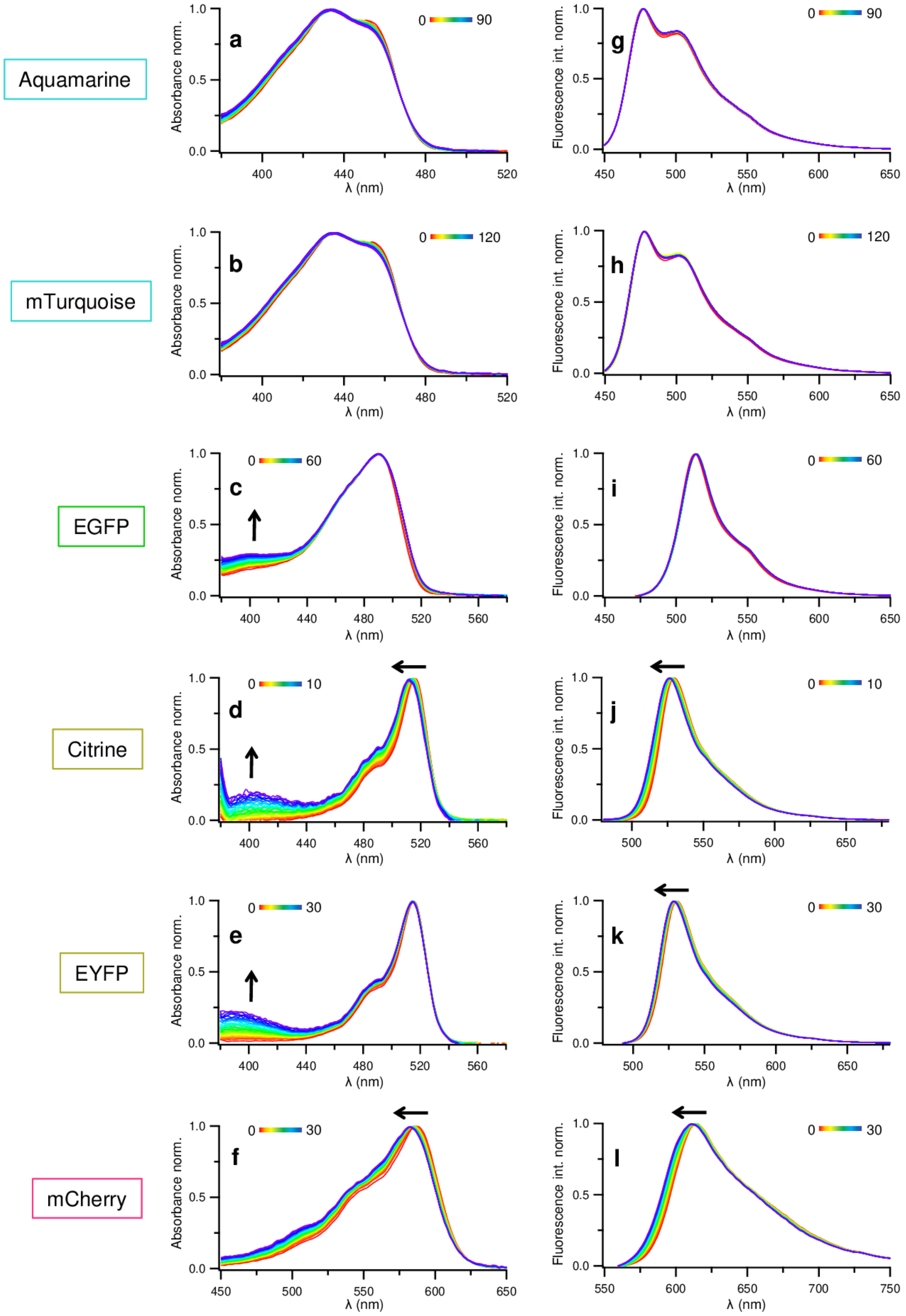

*Figure S13: Time-dependent evolution of normalized absorption spectra (a-f) and normalized fluorescence emission spectra (g-l) during photobleaching using a white light source on BEAM (see Figure S12). The initial maximum absorption of the FP solutions was set to 0.5. Each row presents results obtained for the indicated FP (left).*

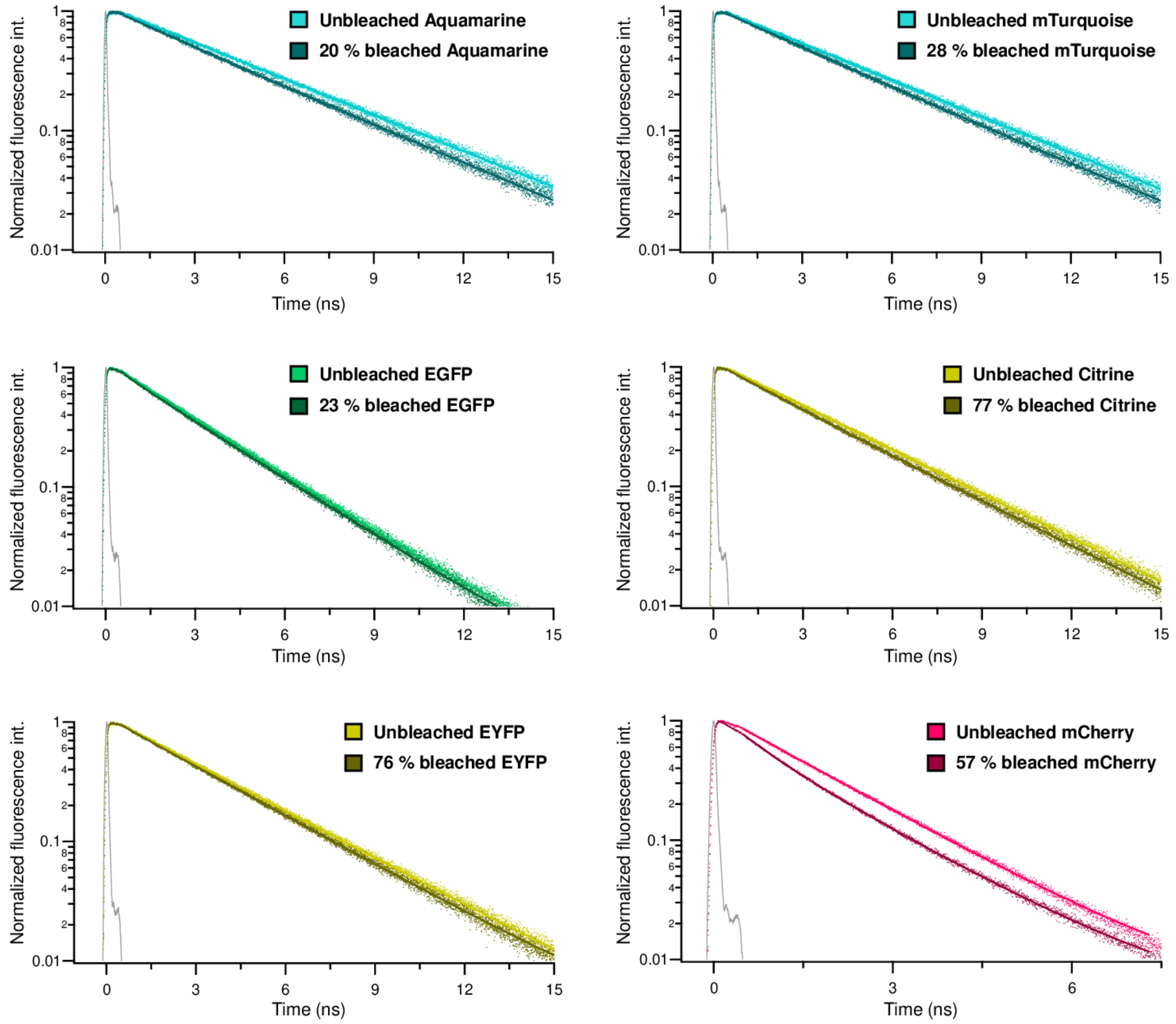

*Figure S14: Representative fluorescence decay curves of* ***purified FPs in solution,*** *recorded before and after photobleaching using a white light source on BEAM (Figure 12). FP solutions were subsequently diluted to maintain an absorbance below 0.1, ensuring accurate measurements of the fluorescence decay curves. Experimental data are shown as dots, and fitted curves as solid lines. A mono-exponential model was used for unbleached FPs, whereas a biexponential model was applied to bleached FPs, with the long lifetime component fixed to the lifetime value measured for the corresponding unbleached FP (see Methods).*

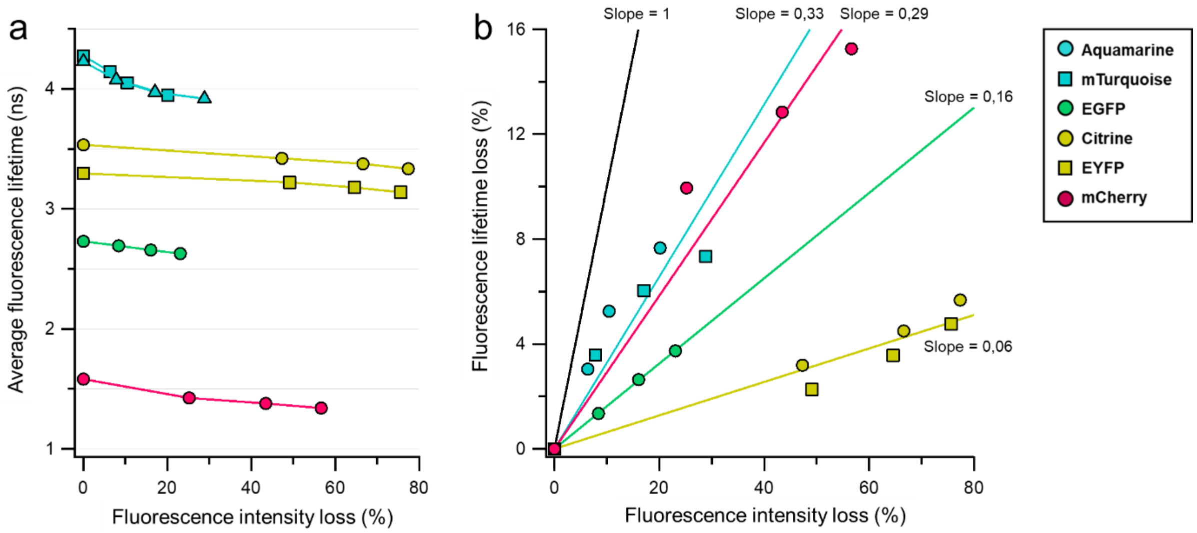

*Figure S15: Evolution of the average fluorescence lifetime of FPs as a function of the relative loss in fluorescence intensity for different FP solutions illuminated using the white light source described in figure S12. Panel (a) shows the average lifetime, and panel (b) shows the lifetime loss. Linear fits obtained by combining all data points for a given color are shown as solid lines. A unit-slope line is included as a visual guide. Representative fluorescence decay curves are shown in Figure S14.*

### E) Model for quantification of photoproducts yields

We developed a model to estimate the relative amounts of the different families of photoproducts identified in FPs solutions after photobleaching. It is assumed that a solution of photobleached FPs contains up to five populations: **unbleached, damaged, dim, colored, and dark** (see Scheme 2). For a given sample, the following measured quantities are required:

- the absorbance of the main band over time, normalized to its initial value: $\boldsymbol{A_{FP,norm}}$
- the absorbance of the band corresponding to the coloured species, normalized to the initial absorbance of the main band: $\boldsymbol{A_{colo,norm}}$
- the fluorescence intensity, normalized to the initial fluorescence intensity: $\boldsymbol{F_{norm}}$
- the fluorescence lifetime, normalized to the initial fluorescence lifetime: $\boldsymbol{\tau_{norm}}$
- the pre-exponential factor of the short-lifetime component from the biexponential fit of the fluorescence decay: $\boldsymbol{\alpha_{damaged}}$

#### E.1 General hypothesis:

We assume that only **unbleached** FP is present initially, at a concentration $[unbleached]_0$, and that a fraction of these proteins converts into four families of photoproducts during illumination. The concentrations of the five species at any given irradiation time are therefore related as:

$$[unbleached]_0 = [unbleached] + [damaged] + [dim] + [colored] + [dark] \qquad \text{(Eq. 1a)}$$

These species can be grouped into fluorescent ("on") and non-fluorescent ("off") populations, Eq. 1d), allowing Eq. 1a to be rewritten as:

$$[unbleached]_0 = [on] + [off] \qquad \text{(Eq. 1b)}$$

with

$$[on] = [unbleached] + [damaged] \qquad \text{(Eq. 1c)}$$

$$[off] = [dim] + [colored] + [dark] \qquad \text{(Eq. 1d)}$$

To obtain quantities more directly relevant to imaging experiments, we also examine the fluorescence signal loss, $\Delta F$, associated with each type of photoproduct

$$\Delta F = F_0 - F = \Delta F_{dark} + \Delta F_{colored} + \Delta F_{dim} + \Delta F_{damaged} \qquad \text{(Eq. 2)}$$

where $F_0$ is the initial fluorescence intensity and $F$ is the fluorescence intensity at the given irradiation time. According to the definitions of Scheme 2 (main text), the unbleached, damaged, and dim species share the same absorption properties. Their combined concentrations can therefore be determined using the Beer–Lambert law:

$$A_{FP} = \varepsilon_{FP,\lambda FP} l([unbleached] + [damaged] + [dim])$$

where $\varepsilon_{FP,\lambda FP}$ is the molar absorption coefficient of the studied FP at its absorption maximum, and $A_{FP}$ is the measured absorbance of the irradiated solution at the same wavelength.

#### E.2 Dark species:

We define the normalized concentration of the dark species as:

$$[dark]_{norm} = \frac{[dark]}{[unbleached]_0}$$

According to equation (1a), the concentration of the dark species can be expressed as:

$$[dark] = [unbleached]_0 - ([unbleached] + [damaged] + [dim]) - [colored]$$

Dividing both sides by $[unbleached]_0$ gives

$$\frac{[dark]}{[unbleached]_0} = 1 - \frac{A_{FP}}{A_{FP,0}} - \frac{[colored]}{[unbleached]_0}$$

With $A_{FP,0}$ the initial absorbance of the FP solution at the wavelength of its absorption maximum.

Finally, we can write:

$$\boldsymbol{[dark]_{norm} = 1 - A_{FP,norm} - [colored]_{norm}}$$

The relative fluorescence loss caused by the formation of dark photoproduct, $BLEACH_{dark}$, is defined as:

$$BLEACH_{dark} = \frac{\Delta F_{dark}}{F_0}$$

Since dark photoproducts do not fluoresce, the fluorescence signal loss is directly proportional to their concentration, yielding:

$$\boldsymbol{BLEACH_{dark} = [dark]_{norm}}$$

#### E.3 Colored species:

We define the normalized concentration of the colored species as

$$[colored]_{norm} = \frac{[colored]}{[unbleached]_0}$$

Due to their absorbance properties, $[colored]$ can be directly determined using the Beer-Lambert law as soon as they are identified with their absorption spectra (position and shape of the band). Nevertheless, the relatively low values of these molar extinction coefficients, which result in weak absorbance bands, can lead to low peak intensities. The latter ones may be significantly influenced by the main absorption band and/or scattering. The contribution from scattering is corrected by baseline subtraction when necessary.

To account for the contribution of the main band, the absorbance of the band corresponding to the colored species must be expressed as

$$A_{colo} = \varepsilon_{FP,\lambda colo} l([unbleached] + [damaged] + [dim]) + \varepsilon_{colo,\lambda colo} l[colored]$$

or equivalently

$$A_{colo} = \varepsilon_{FP,\lambda colo} l \frac{A_{FP}}{\varepsilon_{FP,\lambda FP} l} + \varepsilon_{colo,\lambda colo} l[colored]$$

Where $\varepsilon_{FP,\lambda colo}$ is the molar absorption coefficient of the studied FP, $\varepsilon_{colo,\lambda colo}$ is the molar absorption coefficient of the colored species and $A_{colo}$ is the absorbance of the colored species, all evaluated at the wavelength of the maximum absorption of the colored species.

Solving the concentration of the colored species gives

$$[colored] = \frac{1}{\varepsilon_{colo,\lambda colo} l} A_{colo} - \frac{\varepsilon_{FP,\lambda colo}}{\varepsilon_{FP,\lambda FP} \varepsilon_{colo,\lambda colo} l} A_{FP}$$

which can be rewritten in normalized form as

$$\frac{[colored]}{[unbleached]_0} = \frac{\varepsilon_{FP,\lambda FP}}{\varepsilon_{colo,\lambda colo}} \frac{A_{colo}}{A_{FP0}} - \frac{\varepsilon_{FP,\lambda colo}}{\varepsilon_{colo,\lambda colo}} \frac{A_{FP}}{A_{FP0}}$$

or equivalently

$$\boldsymbol{[colored]_{norm} = \frac{\varepsilon_{FP,\lambda FP}}{\varepsilon_{colo,\lambda colo}} A_{colo,norm} - \frac{\varepsilon_{FP,\lambda colo}}{\varepsilon_{colo,\lambda colo}} A_{FP,norm}}$$

The relative fluorescence loss caused by the formation of these colored photoproducts, $BLEACH_{colored}$ , is defined as

$$BLEACH_{colored} = \frac{\Delta F_{colored}}{F_0}$$

Since colored photoproducts are non-fluorescent, the fluorescence signal loss is directly proportional to their concentration, yielding:

$$\boldsymbol{BLEACH_{colored} = [colored]_{norm}}$$

#### E.4 Dim species:

We define the normalized concentration of the dim species as:

$$[dim]_{norm} = \frac{[dim]}{[unbleached]_0}$$

From Eq.1a, we can write

$$\frac{[dim]}{[unbleached]_0} = \frac{[unbleached]_0 - [dark] - [colored] - [on]}{[unbleached]_0}$$

$$= 1 - [dark]_{norm} - [colored]_{norm} - \frac{[on]}{[unbleached]_0}$$

$$= A_{FP,norm} - \frac{[on]}{[unbleached]_0}$$

The last term can be accessed using the steady state fluorescence intensity during photobleaching[5]:

$$F_{mes} = k\langle f \rangle i_0 (1 - 10^{-A_{FP,on}})$$

where $k$ is a proportionality factor depending on the optical configuration used for the measure, $\langle f \rangle$ is the fluorescence intensity per absorbed photon at the excitation wavelength, $i_0$ is the light intensity at the excitation wavelength, and $A_{FP,on}$ is the absorbance of the emitting species at the wavelength of its absorption maximum. After correction of the inner filter effect using the method described previously[4], we gain access to the extrapolated fluorescence intensity in the linear regime even at high absorbance, which gives us:

$$F = ln(10) k\langle f \rangle i_0 A_{FP,on}$$

Dividing by the initial fluorescence intensity gives

$$\frac{F}{F_0} = \frac{\langle f \rangle A_{FP,on}}{f_0 A_{FP0}}$$

Using the Beer–Lambert law (assuming the damaged species share the same absorption properties as the unbleached species) and assuming $\langle f \rangle$ is proportional to the average fluorescence lifetime $\langle \tau \rangle$ throughout the photobleaching process, we obtain:

$$\frac{F}{F_0} = \frac{\langle \tau \rangle [on]}{\tau_0 [unbleached]_0} <=> \frac{[on]}{[unbleached]_0} = \frac{F_{norm}}{\langle \tau \rangle_{norm}}$$

with $\tau_0$ the initial fluorescence lifetime of unbleached species.

Finally, the normalized concentration of dim species is

$$[\boldsymbol{dim}]_{\boldsymbol{norm}} = \boldsymbol{A}_{\boldsymbol{FP,norm}} - \frac{\boldsymbol{F}_{\boldsymbol{norm}}}{\langle \boldsymbol{\tau} \rangle_{\boldsymbol{norm}}}$$

The relative fluorescence loss caused by the formation of dim photoproducts, $BLEACH_{dim}$ , is defined as

$$BLEACH_{dim} = \frac{\Delta F_{dim}}{F_0}$$

Since the dim photoproducts are non-fluorescent, the loss of fluorescence signal is directly proportional to their concentration, yielding

$$\boldsymbol{BLEACH}_{\boldsymbol{dim}} = [\boldsymbol{dim}]_{\boldsymbol{norm}}$$

#### E.5 Damaged and unbleached species:

Determining the relative populations of damaged and unbleached species follows a similar approach. We define

$$[damaged]_{norm} = \frac{[damaged]}{[unbleached]_0}$$

$$[unbleached]_{norm} = \frac{[unbleached]}{[unbleached]_0}$$

These expressions can be rewritten as

$$\frac{[damaged]}{[unbleached]_0} = \frac{[damaged]}{[on]} \frac{[on]}{[unbleached]_0}$$

$$\frac{[unbleached]}{[unbleached]_0} = \frac{[unbleached]}{[on]} \frac{[on]}{[unbleached]_0}$$

To determine the proportion of damaged or unbleached FPs relative to the total number of photon-emitting FPs, the fluorescence decays of the photobleached solutions are described using a biexponential model. The long lifetime component (fixed to $\tau_0$) corresponds to unbleached species while the short lifetime $\tau_{damaged}$ corresponds to damaged species .

$$I(t) = I_0 \cdot (\alpha_{unbleached} \cdot e^{-t/\tau_0} + \alpha_{damaged} \cdot e^{-t/\tau_{damaged}}) + Bckgd$$

with $I(t)$ the transient fluorescence intensity, $Bckgd$ the background, $I_0$ a proportionality fact, and $\alpha_{unbleached}$ and $\alpha_{damaged}$ are normalized pre-exponential factors such that

$$\alpha_{unbleached} + \alpha_{damaged} = 1.$$

These coefficients directly reflect the relative populations within the emitting species:

$$\frac{[damaged]}{[on]} = \alpha_{damaged},$$

$$\frac{[unbleached]}{[on]} = \alpha_{unbleached}.$$

And finally get:

$$\boldsymbol{[damaged]_{norm} = \alpha_{damaged} \frac{F_{norm}}{\langle\tau\rangle_{norm}},}$$

$$\boldsymbol{[unbleached]_{norm} = \alpha_{unbleached} \frac{F_{norm}}{\langle\tau\rangle_{norm}}.}$$

The fluorescence loss, $BLEACH_{damaged}$, caused by the formation of damaged photoproducts is defined as

$$BLEACH_{damaged} = \frac{\Delta F_{damaged}}{F_0}.$$

Since damaged photoproducts remain partially fluorescent, the corresponding fluorescence loss is not directly proportional to their concentration and must be evaluated separately. Using Eq.2, we can write

$$\frac{\Delta F_{damaged}}{F_0} = \frac{\Delta F - \Delta F_{off}}{F_0}$$

Since all non-fluorescent ("off") species contribute to a fluorescence loss proportional to their concentration, this expression becomes:

$$\frac{\Delta F_{damaged}}{F_0} = \frac{F_0 - F}{F_0} - \frac{[off]}{[unbleached]_0} = 1 - F_{norm} - \left(1 - \frac{[on]}{[unbleached]_0}\right) = \frac{F_{norm}}{\langle\tau\rangle_{norm}} - F_{norm}$$

Finally, we obtain

$$\boldsymbol{BLEACH}_{damaged} = \frac{\boldsymbol{F_{norm}}}{\boldsymbol{\langle\tau\rangle_{norm}}} \boldsymbol{(1 - \langle\tau\rangle_{norm})}$$

#### E.6 Photoproducts quantity and photobleaching fractions:

To obtain values that are independent of the extent of photobleaching, which varies from one FP to another, it is useful to evaluate the contribution of each type of photoproduct relative to the total amount of photoproducts formed. This is achieved by excluding the remaining unbleached FP population from the previously derived expressions.

For a given photoproduct P (dark, colored, dim, or damaged), the **fraction of this photoproduct** can be defined as:

$$FRAC_P = \frac{[P]}{[unbleached]_0 - [unbleached]} = \frac{[P]}{[unbleached]_0} \frac{[unbleached]_0}{[unbleached]_0 - [unbleached]}$$

Using the normalized quantities defined above, this expression becomes

$$\boldsymbol{FRAC_P = \frac{[P]_{norm}}{1 - [unbleached]_{norm}}}$$

Similarly, the **contribution of each type of photoproduct to the total fluorescence loss** can be expressed as

$$FRACB_P = \frac{Flost_P}{F_0 - F}$$

$$<=> \boldsymbol{FRACB_P = \frac{BLEACH_P}{1 - F_{norm}}}$$

### F) Molecular description of FPs photobleaching

#### SDS-Page at high concentration of FP per well (13 µg)

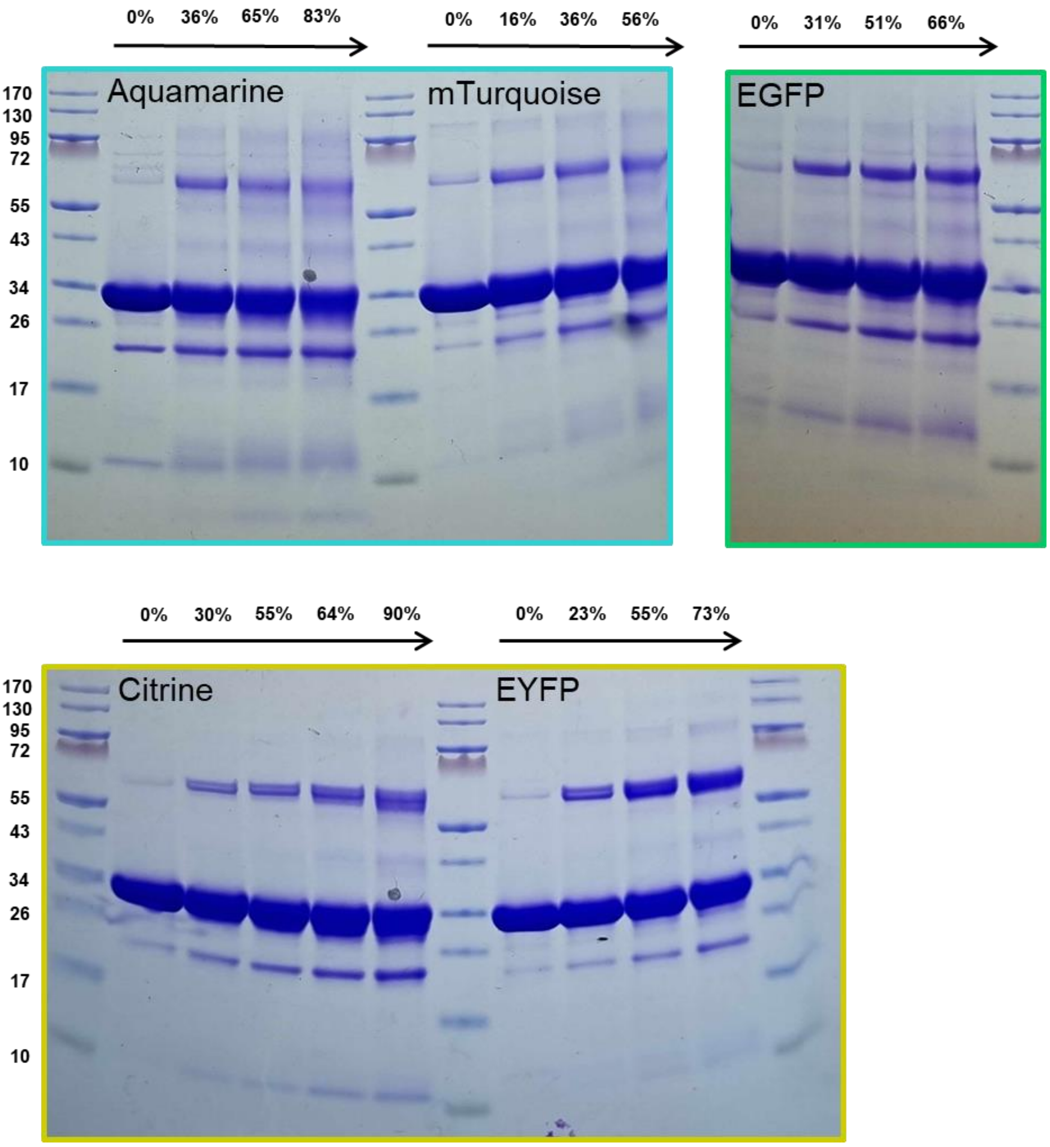

*Figure S17: SDS-PAGE of irradiated FP samples. The molecular mass makers of the protein ladder are indicated in kDa. Each sample was loaded at a concentration sufficient to saturate the main band, enabling visualization of less intense bands corresponding to photoproducts (~ 13 µg of FP per lane). The percentage of fluorescence intensity loss for each sample is indicated above the corresponding lane.* Most non-photobleached FP samples also contain cleaved FP, particularly Aquamarine. This may stem from a similar but less pronounced autolysis reaction.

#### Mass spectrometry under denaturing conditions

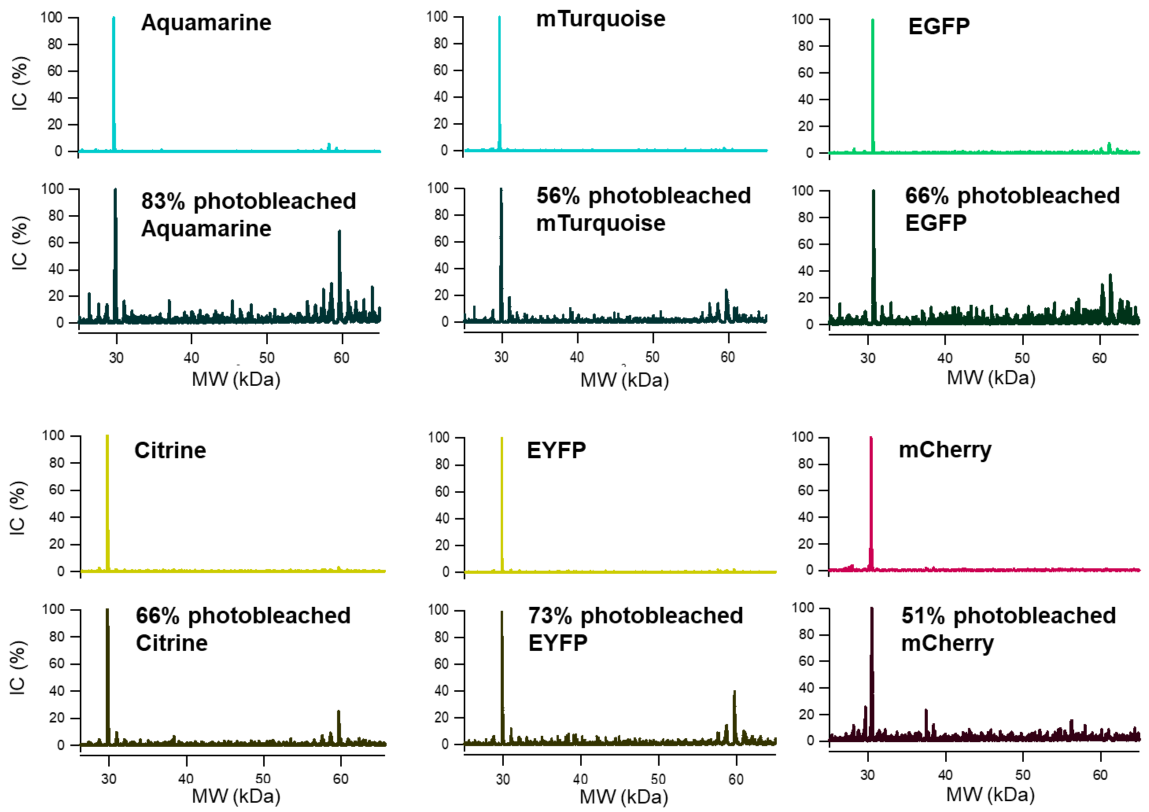

*Figure S18: Evolution of deconvoluted ESI-MS spectra of FPs before and after photobleaching (> 50%).*

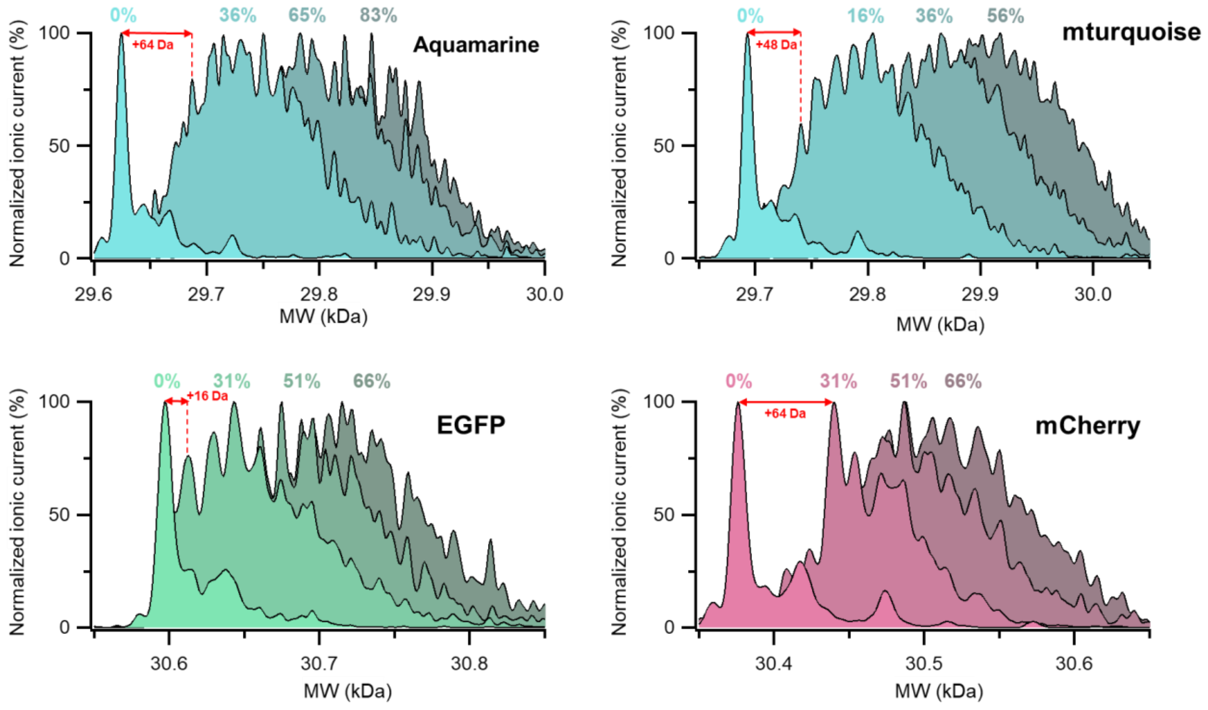

*Figure S19: Evolution of deconvoluted ESI-MS spectra of FPs zoomed in around 30 kDa. The percentage of corresponding fluorescence intensity loss is indicated in the same color. Citrine and EYFP spectra are shown in Figure 6.*

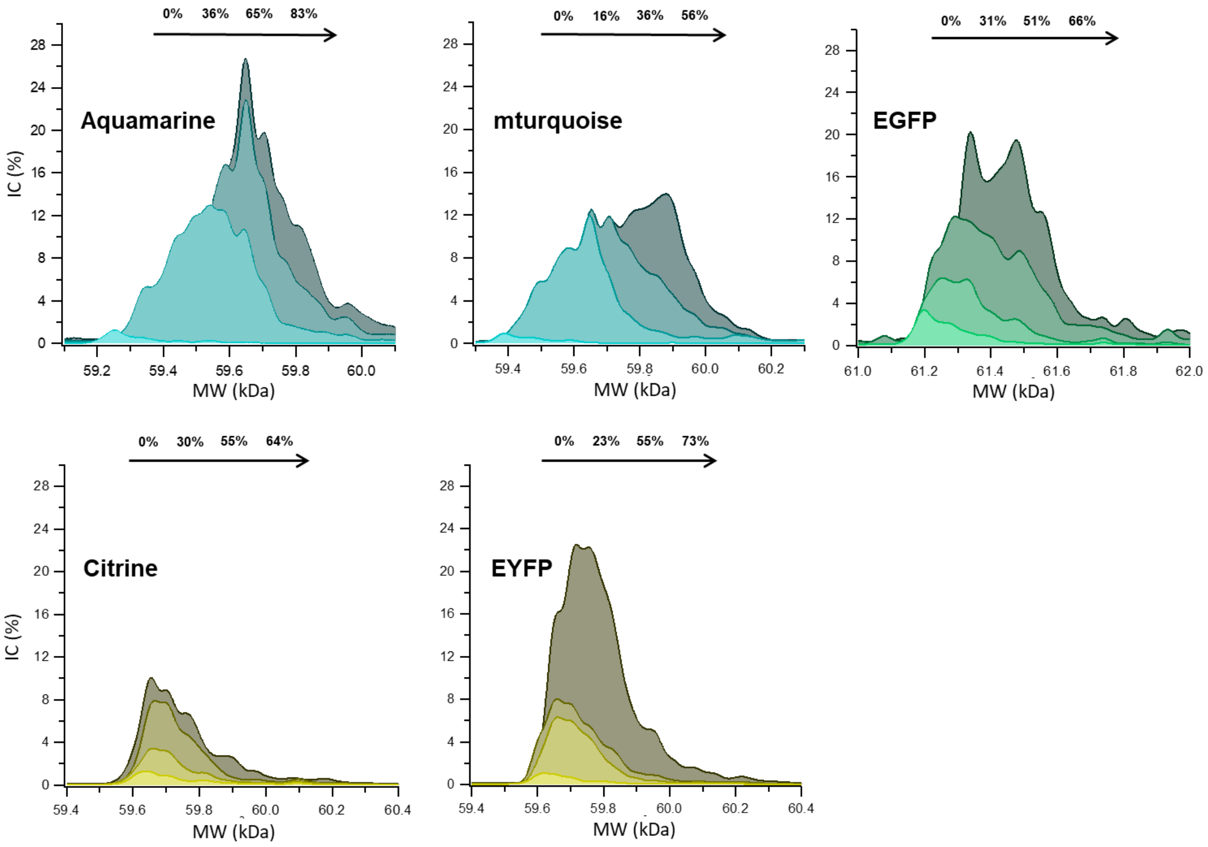

*Figure S20: Evolution of deconvoluted ESI-MS spectra of FPs zoomed in on dimeric species. The percentage of corresponding fluorescence intensity loss is indicated for each spectrum. Due to a low signal-to-noise ratio, spectra were smoothed for clarity.*

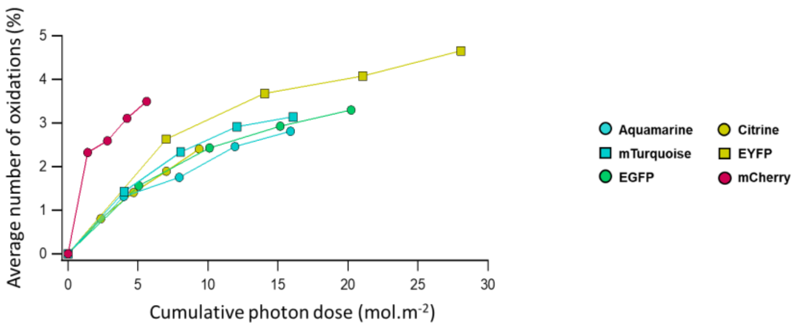

*Figure S21: Evolution of the average number of oxidations per FP as a function of cumulative photon dose, based on the relative peak intensities shown in Figure 6a. The cumulative photon dose was calculated using the equivalent photon flux at the absorption maximum, as indicated in Table S2.*
